# A cryptic plastid and a novel mitochondrial plasmid in *Leucomyxa plasmidifera* gen. and sp. nov. (Ochrophyta) push the frontiers of organellar biology

**DOI:** 10.1101/2022.04.05.487141

**Authors:** Dovilė Barcytė, Karin Jaške, Tomáš Pánek, Tatiana Yurchenko, Tereza Ševčíková, Anežka Eliášová, Marek Eliáš

## Abstract

Complete plastid loss seems to be very rare among secondarily non-photosynthetic eukaryotes. *Leukarachnion* sp. PRA-24, an amoeboid colourless protist related to the photosynthetic algal class Synchromophyceae (Ochrophyta), is a candidate for such a case based on a previous investigation by transmission electron microscopy. Here we characterise this organism in further detail and describe it as *Leucomyxa plasmidifera*, gen. et sp. nov., additionally demonstrating it is the first known representative of a broader clade of non-photosynthetic ochrophytes. We recovered its complete plastid genome, exhibiting a reduced gene set similar to plastomes of other non-photosynthetic ochrophytes yet being even more extreme in sequence divergence. Identification of components of the plastid protein import machinery in the *Leuc. plasmidifera* transcriptome assembly corroborated the organism possesses a cryptic plastid organelle. According to our bioinformatic reconstruction the plastid contains a unique combination of biosynthetic pathways producing haem, a folate precursor, and tocotrienols. As another twist to its organellar biology, *Leuc. plasmidifera* turned out to contain an unusual long insertion in its mitogenome related to a newly discovered mitochondrial plasmid exhibiting unprecedented features in terms of its size and coding capacity. Combined, our work uncovered further striking outcomes of the evolutionary course of semiautonomous organelles in protists.

## 1. Introduction

Despite the benefits conferred to an organism by photosynthesis, this physiological ability has been independently lost by many eukaryote lineages (Hadariová et al. 2018). One such organism, maintained in the American Type Culture Collection (ATCC) as *Leukarachnion* sp. PRA-24, a colourless amoeboid protist that can form reticulate plasmodia, was studied by Grant et al. (2009), who showed it is a stramenopile, specifically a secondarily heterotrophic ochrophyte most closely related to the amoeboid algae *Chlamydomyxa labyrinthuloides* and *Synchroma grande*. This finding raised a possibility that *Leukarachnion* sp. PRA-24 has retained a colourless plastid, but studying the ultrastructure of the protist by transmission electron microscopy (TEM) did not find any candidates for the plastid. No further details on this organism have been published since then, leaving the status of the plastid, and generally the degree of cellular and metabolic reduction related to the loss of photosynthesis, unknown. Furthermore, the taxonomic status of the PRA-24 strain has remained unsettled. Grant et al. (2009) identified it as a potentially novel species putatively related to *Leukarachnion batrachospermi*, a freshwater colourless amoeboid organism forming large anastomosing networks and walled cysts that was described by Lothar Geitler more than 80 years ago (Geitler 1942) but not observed since then. However, morphologically similar heterotrophic reticulate amoebae do not form a phylogenetically coherent group and have evolved many times in different eukaryote lineages (especially in Amoebozoa and Rhizaria; Berney et al. 2015). It thus remains uncertain whether PRA-24 and *Leuk. batrachospermi* are true relatives and whether the former should be placed in the genus *Leukarachnion*.

The PRA-24 strain is not the sole colourless member of the predominantly photosynthetic algal group Ochrophyta. The loss of photosynthesis is a particularly common phenomenon in the class Chrysophyceae, where at least 13 independently evolved non-photosynthetic lineages have been documented so far (Dorrell et al. 2019). Other ochrophyte groups lose photosynthesis less commonly. Two such lineages were described within diatoms (Frankovich et al. 2018; Onyshchenko et al. 2019), at least two separate apochlorotic lineages are known in the class Dictyochophyceae (Sekiguchi et al. 2002; Kayama et al. 2020), and the class Phaeophyceae includes at least one lineage of non-photosynthetic parasites or endophytes of other brown algae (Heesch et al. 2008; Bringloe et al. 2021). In addition, ochrophytes include *Picophagus flagellatus*, a tiny marine heterotrophic flagellate that based on 18S rRNA phylogenies constitutes an independent lineage of its own (Guillou et al. 1999).

Non-photosynthetic ochrophytes span a broad gradient of reduction of photosynthesis-related structures and functions. The chrysophyte *Cornospumella fuschlensis* still retains many plastid-targeted proteins associated with photosystems and light-harvesting complexes, although it lacks enzymes of reductive CO2 fixation (Dorrell et al. 2019). Most other colourless ochrophytes studied in a sufficient detail lack any traces of the photosynthetic machinery, yet still contain reduced plastids (leucoplasts) with their own genome that encode some critical components of specific functional pathways taking place in the organelle (Kamikawa et al. 2015; Kamikawa et al. 2017; Kamikawa et al. 2018; Dorrell et al. 2019; Onyshchenko et al. 2019; Kayama et al. 2020; Bringloe et al. 2021). The set of functional processes retained by the ochrophyte leucoplasts differ between the different independently evolved non-photosynthetic lineages. On the one hand, it may theoretically still include CO2 fixation, as initially suggested for the colourless dictyochophytes *Pteridomonas danica* and *Ciliophrys infusionum* based on identification of the *rbcL* gene encoding the large subunit of the enzyme RuBisCO (Sekiguchi et al. 2002); however, more recent investigations suggested that these *rbcL* sequences might have been contaminants (Kayama et al. 2020). The most extreme reduction has been encountered in the chrysophyte genus *Paraphysomonas*, which contains a vestigial plastid without a genome and hence lacking also the apparatus involved in transcription and translation (Dorrell et al. 2019).

Whether there are any ochrophytes that have secondarily lost the plastid as such is presently unknown. Potential candidates are not only the PRA-24 strain, but also *P. flagellatus*, as no structures identifiable as vestigial plastids have been observed in TEM preparations (Guillou et al. 1999; Grant et al. 2009). Furthermore, no leucoplast could be identified by TEM in *C. infusionum* despite the putative molecular evidence (the *rbcL* gene amplified from the organism) for the presence of a plastid genome (Sekiguchi et al. 2002; but see Kayama et al. 2020), indicating that the apparent absence of the leucoplast at the cytological level must be interpreted with caution. Still, secondarily aplastidic ochrophytes may exist, as evidenced by the recent demonstration that the heterotrophic predatory actinophryids, a subgroup of the traditional protozoan group Heliozoa, constitute a lineage closely related to Ochrophyta and at least one representative, *Actinophrys sol*, exhibits genomic footprints of a photosynthetic ancestry and plastid loss (Azuma et al. 2022).

In this study we set out to test the presence of a cryptic plastid in the PRA-24 strain by molecular approaches. Specifically, we generated transcriptomic and genomic data from the organism and searched for molecular signatures of a plastid. This effort proved fruitful and here we report a complete sequence of a plastid genome of the PRA-24 strain and an *in silico* reconstruction of the major metabolic functions of the elusive plastid organelle beyond any doubts present in this peculiar ochrophyte. As an unexpected twist concerning the organellar biology of the organism, analysis of its mitochondrial genome revealed a large insertion related to a putative mitochondrial plasmid with unprecedented features that we found out to be present in the PRA-24 strain. By exploring environmental DNA (eDNA) data we found out that PRA-24 represents a species that commonly occurs in terrestrial habitats and is part of a broader clade comprised of ecologically similar non-photosynthetic ochrophytes that are yet to be isolated and characterized. Based on a critical evaluation of the historical literature and newly obtained data we formally describe PRA-24 as a new species in a new genus.

Combined, our work has substantially advanced the knowledge of a previously poorly documented non-photosynthetic ochrophyte lineage and further extended the range of unusual organellar biology.

## 2. **Results and Discussion**

### 2.1. *Leukarachnion* sp. PRA-24, redescribed as *Leucomyxa plasmidifera*, gen. et sp. nov., represents a broader ochrophyte clade common in organic-rich terrestrial habitats

The life cycle of the PRA-24 strain studied alternated among vegetative amoeboid cells with different pseudopodial forms, flagellates, and cysts (figure 1). The uninucleate amoebae formed lobopodia (figure 1A,B). Meanwhile, the multinuclear plasmodium formed either filopodia (figure 1C–E) or reticulopodia, resulting in a meroplasmodium. The latter stage was formed only for a very short period of time at the beginning of every sub-culturing. The networks were modest, encompassing only several cell bodies of different sizes. The previous study, on the other hand, reported a large network for the same strain (Grant et al. 2009). Our culturing conditions are likely responsible for the lack of significant anastomosing. Notably, several days after re-inoculation, when bacterial densities increased, cells always became less active. The filopodia-forming plasmodium was very dynamic in terms of its shape and size, with the main contracted body measuring up to 15 µm in length. The smallest measured cell body was 1.5 µm in length. The filopodia were of different thickness with the finest threads usually stretching out posteriorly and laterally from the bulging ones directly connected to the main body. In our cultures, one to six, mostly two or three, filopodia radiated from the main body (figure 1C–E). The active movement of the main body and pseudopodia was observed, including building, stretching and pulling in the filopodia and reticulopodia, resulting in an actively migrating plasmodium. Little bulbous protrusions could be seen to move inside the threads towards the main body or in the opposite direction.

**Figure 1.**
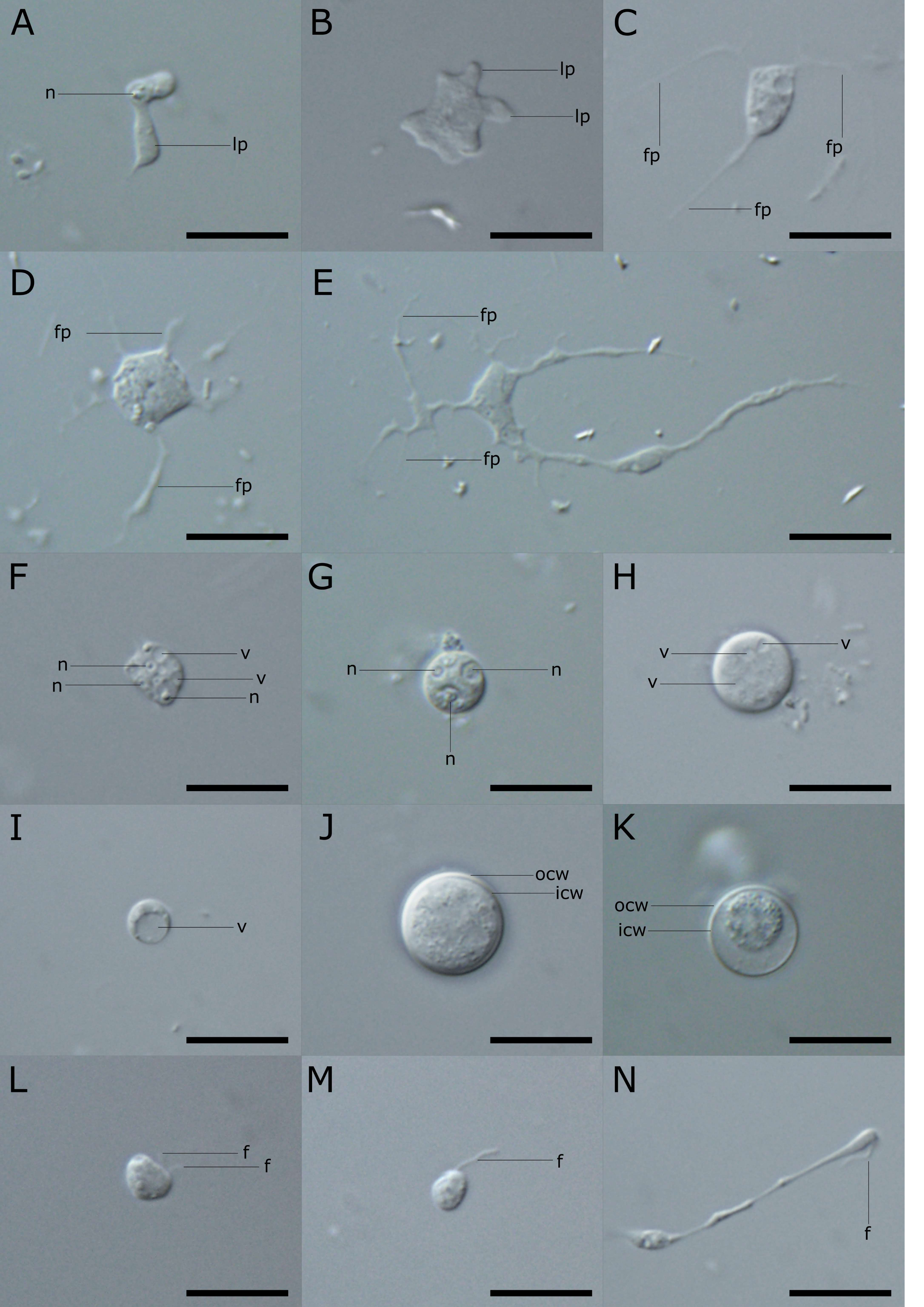
Light microscopy of different life stages of *Leucomyxa plasmidifera* sp. nov. (*Leukarachnion* sp. PRA-24). (A–B) Uninucleate amoebae with lobopodia (lp). (C–D) Multinucleate plasmodia with different number of filopodia (fp). (F–H) Resting stages with prominent nuclei (n) and vacuoles (v). (I) Cell with a single huge vacuole. (J–K) Cysts with a doubled-layered thick covering: inner (icw) and outer (ocw) cell wall. (L) Biflagellate cell with subapical flagella (f). (M) Uniflagellate cell with an apical flagellum. (N) Interaction of two cells. Scale bars: 10 µm.

Although the amoeboid form is most characteristic for PRA-24, our cultures were dominated by resting stages, i.e. irregularly roundish cells with a slight movement or none at all. Apart from nuclei with prominent nucleoli, they also contained numerous randomly distributed vacuoles (figure 1F–H). We have also observed cells containing a single huge vacuole (figure 1I). The same cell type was also found in the related amoeboid alga *S. grande* (Synchomophyceae, Ochrophyta), and considered an intermediate stage between migrating and sessile amoebae (Koch et al. 2010). During the several weeks of cultivation, almost all PRA-24 cells transformed into cysts. They could be distinguished from the vegetative cells by having a regular spherical shape and being covered by a thick double-layered cell wall (figure 1J,K). Their size varied from 2.5–7.0 µm in diameter (n=90). This size range generally matched the one of vegetative non-branching cells, suggesting that every cell is capable of encystation when growth conditions are no longer favourable. The cell wall of the cysts was smooth and did not contain any obvious protrusions or germination pores (figure 1J,K).

Sometimes, the cell content inside the cyst was considerably shrunken but appeared to be active (figure 1K). Such coated cells were possibly undergoing mitotic division.

In addition to active amoeboid cells, pre-cysts and cysts, we have also observed two types of flagellated cells (figure 1L,M). This is in agreement with Grant et al. (2009), who noted the presence of flagellates in their PRA-24 cultures but they did not provide any details. The first type of observed swimming cells had two flagella of unequal length. They were inserted subapically (figure 1L). The biflagellate cells ranged from oval to bean-shaped in shape and were approximately 3.0 µm long and 2.5 µm in wide. They moved fast and rotated while swimming. Such cells likely represent zoospores. The second type of motile cells included potentially uniflagellate cells with a flagellum being inserted apically (and a posterior flagellum, if present, not discernible; figure 1M). Their size reached up to 3.0 µm in length and 2.0 µm in width. They could possibly represent gametes. This hypothesis is also supported by the fact that we did observe a peculiar interaction of two such cells with one of them exhibiting an active single-flagellum movement and dragging the second one while it slowly crawls towards the swimming one (figure 1N). However, the elucidation of a full life cycle of the PRA-24 strain, including the alternation of different morphotypes, awaits further detailed investigations.

To complement the previous cursory investigation of the PRA-24 strain ultrastructure by Grant et al. (2009), we carried out our own TEM study of the organism; see supplementary electronic material, figure S1 (representative images) and note S1 (a detailed discussion). Briefly, or findings concerning the amoeboid cells and cysts correspond well with the account by Grant at al. Notably, our TEM investigation confirmed the presence of flagellated cells, with two flagella and their basal bodies detected (figure S1G–I). Though not complete, the observed structures of *Leuc. plasmidifera* flagellar apparatus still make it possible to consider their homology with other ochrophyte representatives. The basal bodies of the two flagella were roughly perpendicular to each other and overlapped their proximal ends in a clockwise orientation (figure S1H). Such configuration resembles that of, for example, chrysophytes (Andersen 1990). A transition (proximal) helix with four gyres above the transition (basal) plate was detected in the anterior flagellum. In addition, a dense band linking the axonemal doublets to an unidentified structure was also present (figure S1I). In various representatives from closely related classes (including, for example, *Spumella* spp., Chrysophyceae), four gyres are quite common and the aforementioned dense band connects axonemal doublets to the plasmalemma (Hibberd 1979). Although a more detailed reconstruction of the ultrastructure of the *Leuc. plasmidifera* flagellar apparatus would be desirable, the results of our TEM observations are consistent with *Leuc. plasmidifera* representing a lineage affiliated with Chrysophyceae.

We used the 18S rRNA gene to update the understanding of the phylogenetic position of the PRA-24 strain. The PRA-24 sequence as determined in this study by genome and transcriptome sequencing (see below) differs from the previously published PCR-based sequence (FJ356265.2) at its 5’ end, obviously due to the latter being chimeric (the differing 5’ segment most likely derived from a fungus, judging from a sequence similarity search). By exploring the non-redundant (nr) nucleotide sequence database and metagenomic assemblies at the National Center for Biotechnology Information (NCBI) we identified 51 complete or partial 18S rRNA gene sequences (excluding those recognized as chimeras) that in a phylogenetic tree formed a highly supported clade (94% of bootstrap proportions) with the respective PRA-24 sequence to the exclusion of sequences from cultured, taxonomically identified organisms (figure 2, supplementary electronic material, table S1). The environmental sequences in this clade exhibited substantial disparity, differing from the (updated) PRA-24 sequence at as much as 6% of positions compared (with the lowest similarity values observed for partial sequences matching the most variable regions of the 18S rRNA gene). In accord with previous analyses (Schmidt et al. 2015), the clade comprised of PRA-24 and the related eDNA sequences was further united into a higher-order fully supported grouping with two additional clades, one comprised of multiple sequences attributed to *Chlamydomyxa labyrinthuloides* and the other containing sequences from the genus *Synchroma* and its specific relatives (*Guanchochroma wildpretii* and *Chrysopodocystis socialis*). This large grouping in turn formed a clade sister to the class Chrysophyceae.

**Figure 2.**
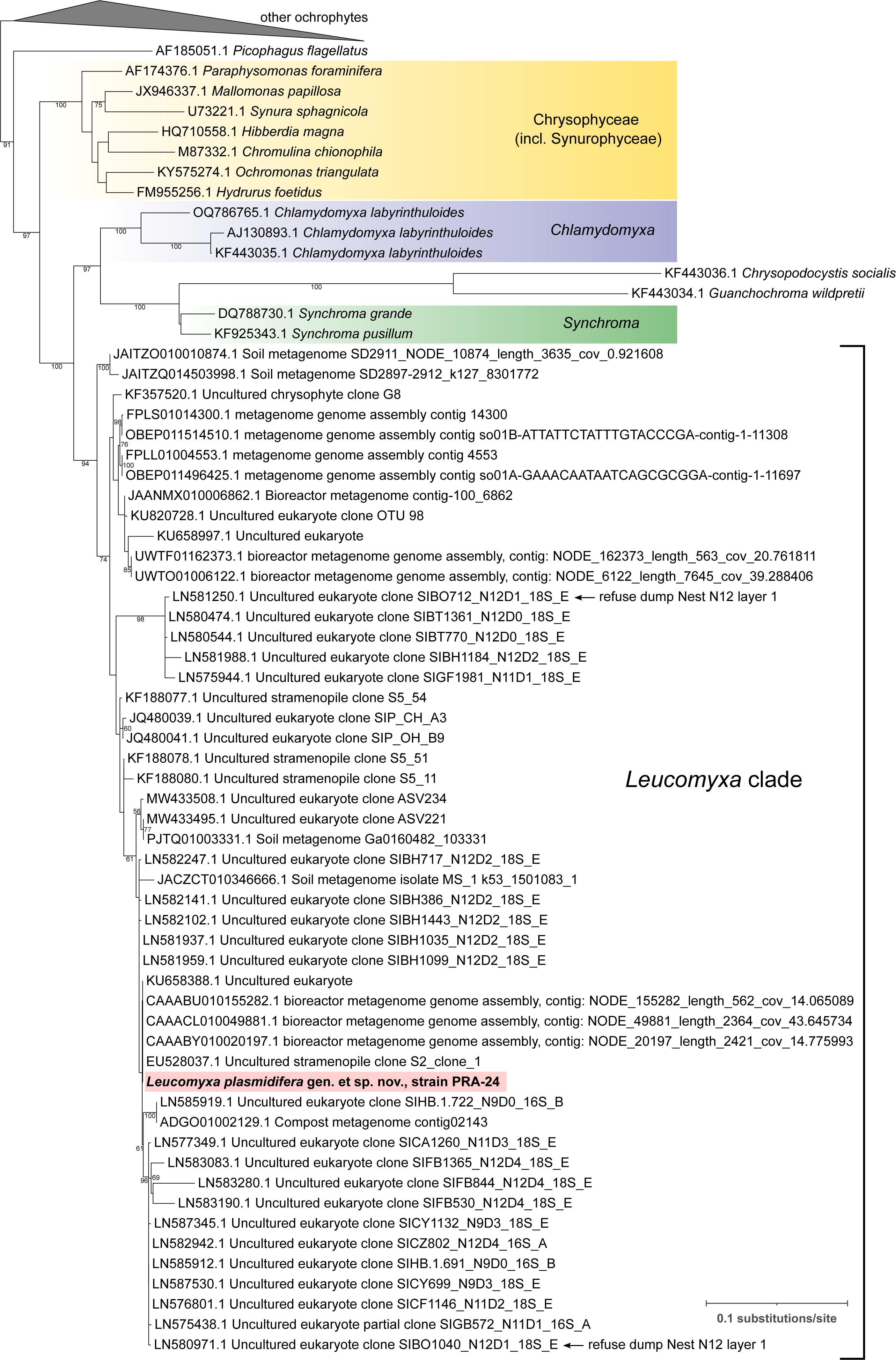
The phylogenetic position of *Leucomyxa plasmidifera* PRA-24 based on the 18S rRNA gene. The tree was inferred from a multiple nucleotide sequence alignment (1,729 positions) using IQ-TREE (TIM2+F+I+R5 substitution model selected by the program as the best fitting to the data). The numbers at branches represent bootstrap support values (shown when >50%). In addition to the updated 18S rRNA gene sequence from *Leuc. plasmidifera* PRA-24 (see main text), included were 49 non-chimeric sequences from environmental surveys (both amplicon-based and metagenomic) that clustered with the *Leuc. plasmidifera* sequence in preliminary analyses (details on the sequences are provided in supplementary electronic material, table S1) and sequences from selected representatives of all known ochrophyte classes and lineages *incertae sedis*. For simplicity, only a part of the tree including taxa of special interest (those in the phylogenetic neighbourhood of *Leuc. plasmidifera*) is shown in full; the rest (other ochrophytes) was collapsed as a triangle on the top of the tree. A full tree is provided in the Newick format in supplementary electronic material. The two sequences marked with the note “refuse dump Nest N12 layer 1” come from a particular environmental sample (a refuse dump created by a leaf-cutter ant) from which sequences specifically related to the 16S rRNA gene from the *Leuc. plasmidifera* PRA-24 plastid were reported (supplementary electronic material, figure S4).

These results indicate that the PRA-24 clade represents a broader separate taxon, at least one genus with multiple species. The eDNA data additionally provide an important insight into the ecology of this emerging ochrophyte group. The PRA-24 strain was isolated from a salt march in Virginia, but details on the original habitat and locality were not reported by Grant et al. (2009) and are not available from the metadata provided by ATCC. To illuminate the ecological range of the respective organism, we extracted from its genome assembly the sequence of the highly variable rRNA ITS2 region and explored the occurrence of this “barcode” in raw metagenomic data using the PebbleScout search tool (Shiryev and Agarwala 2023). 461 metagenomic samples were retrieved matching the query with the maximal PBSscore value (i.e. 100) and 100% coverage (supplementary electronic material, table S2), indicating the presence of an identical or highly similar sequence in the metagenome. In terms of their origin, these metagenomes predominantly came from different types of soil samples and similar habitats (rhizosphere, earthworm microbiome etc.) taken at different places around the globe, indicating the species represented by PRA-24 is a common terrestrial organism not specifically associated with salty spots. The PRA-24 relatives detected on the basis of environmental 18S rRNA gene sequences come from a similar range of habitats, including soil, rhizosphere, compost, refuse dumps created by leaf-cutter ants (*Atta colombica*), a lake sediment, and even in human-made habitats such as bioreactors and wastewater treatment plants (supplementary electronic material, table S1), which indicates that the whole PRA-24 clade embraces ecologically similar organisms that tend to thrive in organic-rich terrestrial environments. Given the general morphology of the PRA-24 strain itself as well as its closest known relatives mentioned above, the eDNA sequences specifically affiliated to PRA-24 are expected to be derived from similar amoeboid organisms, presumably also non-photosynthetic based on evidence presented below.

The question now arises if the PRA-24 clade includes any previously formally described taxon, above all *Leuk. batrachospermi* considered a putative PRA-24 relative by Grant et al. (2009). The original description by Geitler (1942) is the sole account on *Leuk. batrachospermi* we could find in the literature; its summary and Geitler’s later investigation of a similar unidentified organism is provided in supplementary electronic material, note S2. Morphologically, both *Leuk. batrachospermi* and the PRA-24 strain represent a phylogenetically highly heterogeneous category of reticulopodial heterotrophic “rhizopods” whose phylogenetic relationships cannot be discerned without molecular data (Berney et al. 2015). From this perspective, the identification of the PRA-24 strain as a member of the genus *Leukarachnion* by Grant et al. (2009) must be taken with caution and further scrutinized. In fact, the authors considered a specific relationship of PRA-24 to *Leuk. batrachospermi* only on the account of both forming a generically similar plasmodial network (meroplasmodium) and cysts. However, they also noted PRA-24 being much smaller than *Leuk. batrachospermi*. The size difference concerns also the cysts, in PRA-24 being (in their mature form) twice as small as reported for *Leuk. batrachospermi* (10–14 µm). Furthermore, the cysts as of *Leuk. batrachospermi* as depicted and described by Geitler (1942) differ from the cysts of PRA-24 by containing stubby or spiky protrusions outside the cyst wall, while cysts of PRA-24 do not form such structures. Even more significant for the assessment of the potential relationship of the PRA-24 strain and *L. batrachospermi* seem to the fact that the former produces flagellated stages whereas Geitler (1942) did not observed any flagellates for *Leuk. batrachospermi*, and that PRA-24 is bacteriovorous, making its nutritional strategy very different from the highly specialized feeding of red algal reproductive cells documented for *Leuk. batrachospermi*.

Combined, there is no doubt that PRA-24 and *Leuk. batrachospermi* are different species, and we additionally argue the case is weak for presupposing that *Leuk. batrachospermi* belongs to a closer phylogenetic neighbourhood of PRA-24. As mentioned above, eDNA documents the existence of a broader ochrophyte clade comprising PRA-24 together with uncultured organisms recorded in terrestrial habitat types rich in organic matter and bacteria, which is consistent with a notion that they are all nutritionally similar to the PRA-24 strain, i.e. relying on bacteriovory (with possible contribution of osmotrophy, untested for PRA-24); at any rate, eDNA evidence for PRA-24 relatives occurring in freshwater habitats, which would be expected if *L. batrachospermi* was related to PRA-24, was not seen. We thus consider it is safe to conclude that the PRA-24 strain is not a new species of the presently monotypic genus *Leukarachnion*. The specific features of *Leuk. batrachospermi* in fact suggest its phylogenetic affinity to Vampyrellida, an unrelated amoeboid group in the supergroup Rhizaria, and there seems to be no other previously established genus of reticulopodial amoeboid protists that would be a good candidate for the taxonomic home of PRA-24 (additional details are discussed in supplementary electronic material, note S2). Hence, to taxonomically accommodate the PRA-24 strain we describe it as a new species in a new genus, *Leucomyxa plasmidifera* (the formal taxonomic treatment is provided below in section 4). Furthermore, as there are no candidates for previously described specific relatives of *Leuc. plasmidifera*, and given its deep phylogenetic separation in the 18S rRNA gene tree (figure 2) from the existing orders Synchromales (comprising the genus *Synchroma*, with *Chrysopodocystis* and *Guanchochroma* apparently also falling into it) and Chlamydomyxales (containing the genus *Chlamydomyxa*), we formally propose in section 4 a new family, Leucomyxaceae, and a new order, Leucomyxales (presumably corresponding to the *Leucomyxa* clade as delimited by the 18S rRNA gene phylogeny), to provide a proper classification of *Leuc. plasmidifera* in the hierarchical Linnean system. We leave the class-level assignment of *Leuc. plasmidifera* open until its phylogenetic position is studied by phylogenomics (work in progress).

### 2.2. A highly reduced plastid genome occurs in *Leucomyxa plasmidifera*

As already mentioned, previous investigations by TEM did not reveal any candidate structures that might be identified as a plastid in *Leuc. plasmidifera* (Grant et al. 2009). Our own TEM study remained inconclusive as to the identification of structures that could be unambiguously identified as putative plastids (leucoplasts) in *Leuc. plasmidifera* cells (supplementary electronic material, note S1). To address the question of the plastid status in *Leuc. plasmidifera* by an alternative approach, we used the Illumina HiSeq sequencing technology to perform a survey sequencing of a DNA sample isolated from the *Leuc. plasmidifera* culture. The longest scaffolds in the resulting sequence assembly were derived from bacteria, whereas the *Leuc. plasmidifera* nuclear genome was highly fragmented due to low read coverage (∼2×). Hence, to create a resource for exploration of the gene repertoire of the organism we also sequenced the transcriptome of *Leuc. plasmidifera*. The transcriptome assembly contained 19,283 contigs and seems to be highly representative of the expected *Leuc. plasmidifera* gene complement, with all genes in a stramenopile-specific reference BUSCO genes set represented as complete sequences (supplementary electronic material, figure S2). 40% of them were found to be present in more than one copy, and indeed alterative transcript variants for many individual genes were identified during manual analyses, reflecting alternative splicing variants or different alleles. Manual analyses of the transcriptome assembly additionally revealed a minor bacterial contamination (derived primarily from a *Vibrio* species), which was taken into account in further analyses.

Strikingly, searching the *Leuc. plasmidifera* DNA sequence contigs with reference plastid genes retrieved three contigs that resembled parts of a plastid genome lacking photosynthesis-related genes. Owing to overlaps the three contigs could be manually joint to obtain a single circular-mapping sequence of 42,214 bp that exhibits the conventional plastid genome architecture with two inverted repeats separated by the long and short single-copy regions (figure 3A). We refer to the assembled ∼42 kbp sequence as to the *Leuc. plasmidifera* plastome, which is further corroborated by phylogenetic analyses presented below.

**Figure 3.**
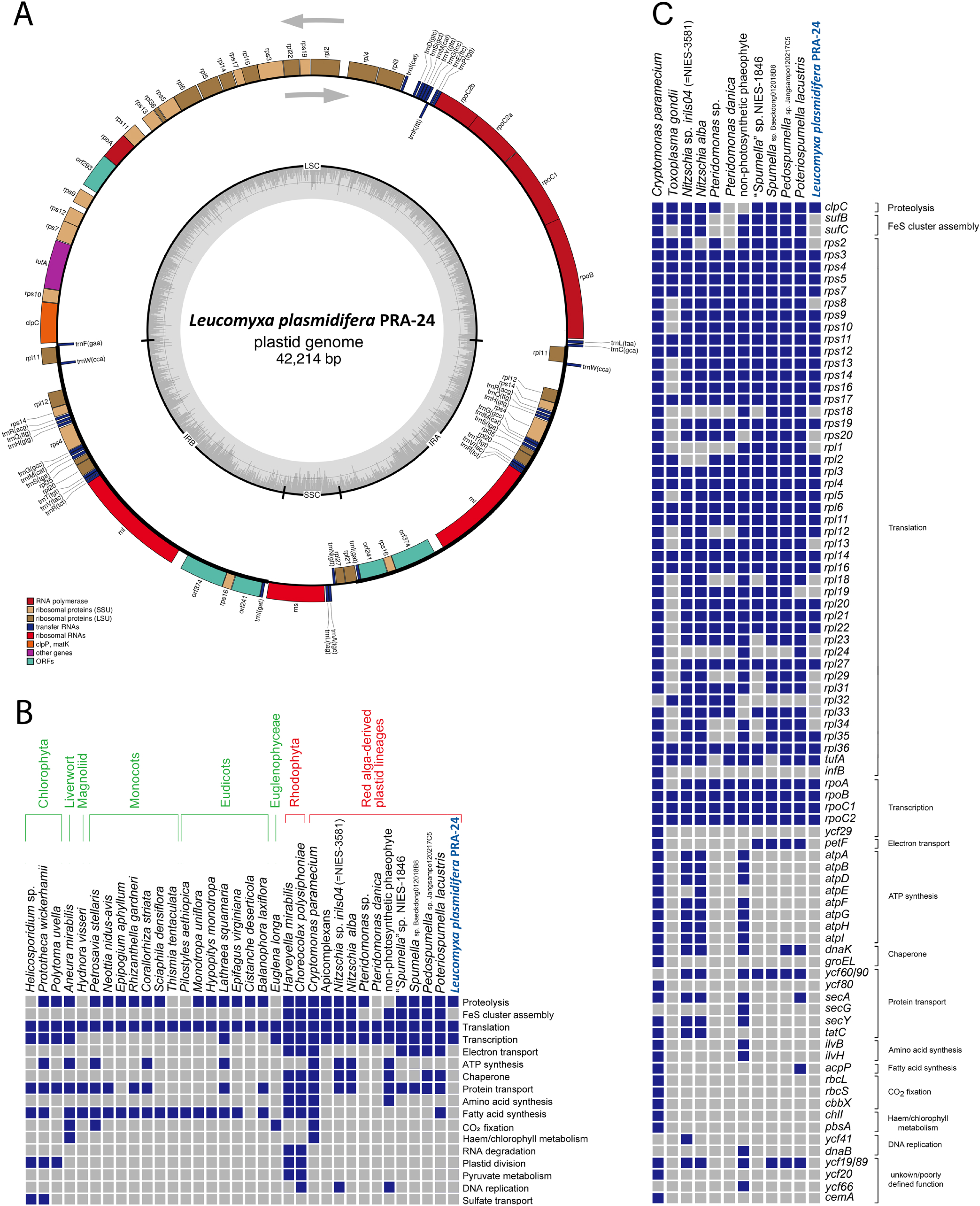
Plastid genome of *Leucomyxa plasmidifera*. (A) Map of the genome. Genes are shown as boxes (coloured according to the functional category they belong to; see the graphical legend) facing inward or outward of the circle depending on the direction in which they are transcribed (clockwise or counter-clockwise, respectively; see the grey arrows). The thickenings of the outer circle delimit regions representing inverted repeats. The internal circle in grey indicates the variation in the GC content along the plastid genome. LSC – long single-copy region; SSC – short single-copy region; IRA and IRB – inverted repeats. (B) Functional categories represented in the gene sets of plastid genomes of diverse non- photosynthetic algae and plants, including *Leuc. plasmidifera*. (C) Occurrence of protein- coding genes in plastid genomes of non-photosynthetic ochrophytes and selected other eukaryotes with a colourless plastid of a red-algal origin. Parts B and C were adopted with modifications (addition and reordering of taxa) from Kayama et al. (2020).

Disregarding the duplications due to the presence of the inverted repeats, we identified in the plastome 38 protein-coding genes (including three unidentified ORFs), 26 tRNA genes, and two rRNA genes (16S and 23S). Only the 23S rRNA gene is present in the inverted repeat in the *Leuc. plasmidifera* plastid genome, whereas the 16S rRNA gene is located in a single copy in the short single-copy region (figure 3A). An analysis of the *Leuc. plasmidifera* plastome with FACIL (Dutilh et al. 2011) did not indicate any signal for any codon having a non-standard meaning (supplementary electronic material, figure S3A), which agrees with the fact that the plastome contains a standard set of tRNA genes directly comparable to tRNA gene sets in plastomes of other ochrophytes (supplementary electronic material, table S3).

A comparison of the repertoire of biological functions mediated by protein-coding genes in the plastomes of *Leuc. plasmidifera* and selected other non-photosynthetic eukaryotes is shown in figure 3B, a comparison of the occurrence of protein-coding plastid genes in *Leuc. plasmidifera* and a smaller subset of non-photosynthetic taxa is provided in figure 3C, and a detailed look at the gene content of plastid genomes of *Leuc. plasmidifera* and diverse ochrophytes is provided in supplementary electronic material, table S3. The *Leuc. plasmidifera* plastid genome lacks all genes encoding components of photosystems and the electron-transport chain, the respective biogenesis factors, and chlorophyll-synthesis enzymes, indicating complete loss of photosynthetic functions. In contrast to some other non-photosynthetic plastids (Hadariová et al. 2018; Bringloe et al. 2021; Pánek et al. 2022), it also lacks genes for the FoF1 ATP synthase, RuBisCO subunits, and proteins involved in RuBisCO expression and regulation. Lost are also all other plastid genes that in other ochrophytes encode enzymes of plastid-localised biosynthetic pathways, including the synthesis of fatty acids, branch amino acids, and thiamin. Further missing are genes for components of the membrane protein-targeting systems SEC and TAT, and for the chaperones DnaK and GroEL. A salient feature of the *Leuc. plasmidifera* plastid genome is the absence of the genes *sufB* and *sufC* encoding the critical components of the SUF system of the iron-sulfur (Fe-S) cluster assembly. Both *sufB* and *sufC* are retained by plastid genomes of virtually all ochrophytes, including the non-photosynthetic ones (figure 3C; Kamikawa et al. 2018; Dorrell et al. 2019; Bringloe et al. 2021). Both are also kept in the plastid genome of parasitic colourless red algae (Salomaki et al. 2015; Preuss et al. 2020) and non-photosynthetic cryptophytes (Tanifuji et al. 2020), and *sufB* is present in plastid genomes of apicomplexans (except for *Plasmodium*) and some other non-photosynthetic myzozoans (Mathur et al. 2019). The only other exception among ochrophytes is the dictyochophyte genus *Pteridomonas* (Kayama et al. 2020) lacking both *sufB* and *sufC* like *Leuc. plasmidifera* (figures 2B and 2C).

Thus, the products of the genes retained in the *Leuc. plasmidifera* plastome seem to belong to only three functional categories: transcription (RNA polymerase subunits), translation (ribosomal proteins, the translation elongation factor Tu, rRNAs and tRNAs), and protein turnover (ClpC). However, this list may be incomplete, as the function of three putative protein-coding genes could not be deduced because of the lack of discernible homology to other proteins even when using the highly sensitive homology detection tools HHpred and Phyre2. No insights into the origin of these putative genes were obtained even when the tertiary structure of their protein products was predicted with AlphaFold2 and the structural models compared with FoldSeek to a broad database of experimentally determined and predicted proteins structures. One of these unidentified ORFs, *orf293*, is localised downstream of *rpoA* that in other plastid genomes is occupied by the *rpl13* gene. No *rpl13* is recognizable elsewhere in the *Leuc. plasmidifera* plastid genome, so we visually inspected a multiple alignment of Orf293 and ochrophyte Rpl13 proteins, but the former really does not seem to fit the sequence pattern of Rpl13; if *orf293* is an *rpl13* ortholog, it must have diverged extremely. The other two, *orf374* and *orf241*, are present in the inverted repeat region and flank the gene *rps16*. The putative proteins encoded by these ORFs include two predicted transmembrane helices each (being thus the only transmembrane proteins encoded by the whole plastid genome). We tried to match them to plastid proteins with a similar general architecture or those encoded by genes with an analogous position in the genome, but without finding any plausible candidates for homologs; the origin of these proteins and their function thus remain unknown.

No 5S rRNA gene is discernible in the assembled *Leuc. plasmidifera* plastome sequence. There is, in fact, an unannotated region present in the genome directly downstream of the 23S rRNA gene, exactly where the 5S rRNA gene is expected to be located. However, no match of this region, or any other in the *Leuc. plasmidifera* plastid genome, to 5S rRNA was detected even when we employed the most sensitive approach available, utilizing covariance models built for alternative 5S rRNA arrangement including a circularly permuted structure of the RNA (such as was described for the mitochondrial 5S rRNA in brown algae; Valach et al. 2014). Interestingly, no 5S rRNA gene was annotated in the plastid genomes certain other non-photosynthetic ochrophytes, namely the chrysophyte “*Spumella*” sp. NIES-1846 (Dorrell et al. 2019) and the dictyochophytes of the genus *Pteridomonas* (Kayama et al. 2020).

Whether this means real loss of the 5S rRNA gene and thus presumably the 5S rRNA molecule as such from the plastidial ribosome of this species remains an open question. An alternative possibility, consistent with evidence for extremely rapid plastid genome evolution in the *Leuc. plasmidifera* presented in the next section, is that a 5S rRNA gene is present but has diverged beyond recognition with the currently available computational approaches.

### 2.3. Plastid genes in *Leucomyxa plasmidifera* are extremely divergent

To formally verify the presupposed origin of the circular-mapping plastome-like sequence described above, we included 30 proteins encoded by this putative plastid genome in a concatenated alignment together with the respective plastome-encoded orthologs from a reference set of ochrophyte species (selected such as to have represented all classes with plastid genome data available) and several non-ochrophyte algae treated as an outgroup (supplementary electronic material, table S4). The supermatrix (6,627 aligned amino acid positions) was analysed with the maximum likelihood (ML) method and the complex model LG+C60+F+G. The resulting tree (figure 4) is generally congruent with recent phylogenetic analyses based on multiple plastid genome-encoded proteins (Ševčíková et al. 2019; Han et al. 2019; Kim et al. 2020; Barcytė et al. 2021; Di Franco et al. 2022; Barcytė et al. 2022). We note that we could not include in the analysis the closest relatives of *Leuc. plasmidifera* as identified by nuclear genes-based phylogenies, such as species of the genera *Chlamydomyxa* or *Synchroma*, since plastid genome sequences have not yet been reported from them and a previously published transcriptome assembly from *Synchroma pusillum* (Keeling et al. 2014) does not include protein-encoding plastidial transcripts corresponding to the genes retained in the *Leuc. plasmidifera* plastome. In the absence of these organisms the plastid genome of *Leuc. plasmidifera* is expected to place this organism as a sister lineage of the class Chrysophyceae, which is exactly what is observed in our tree (figure 4).

**Figure 4.**
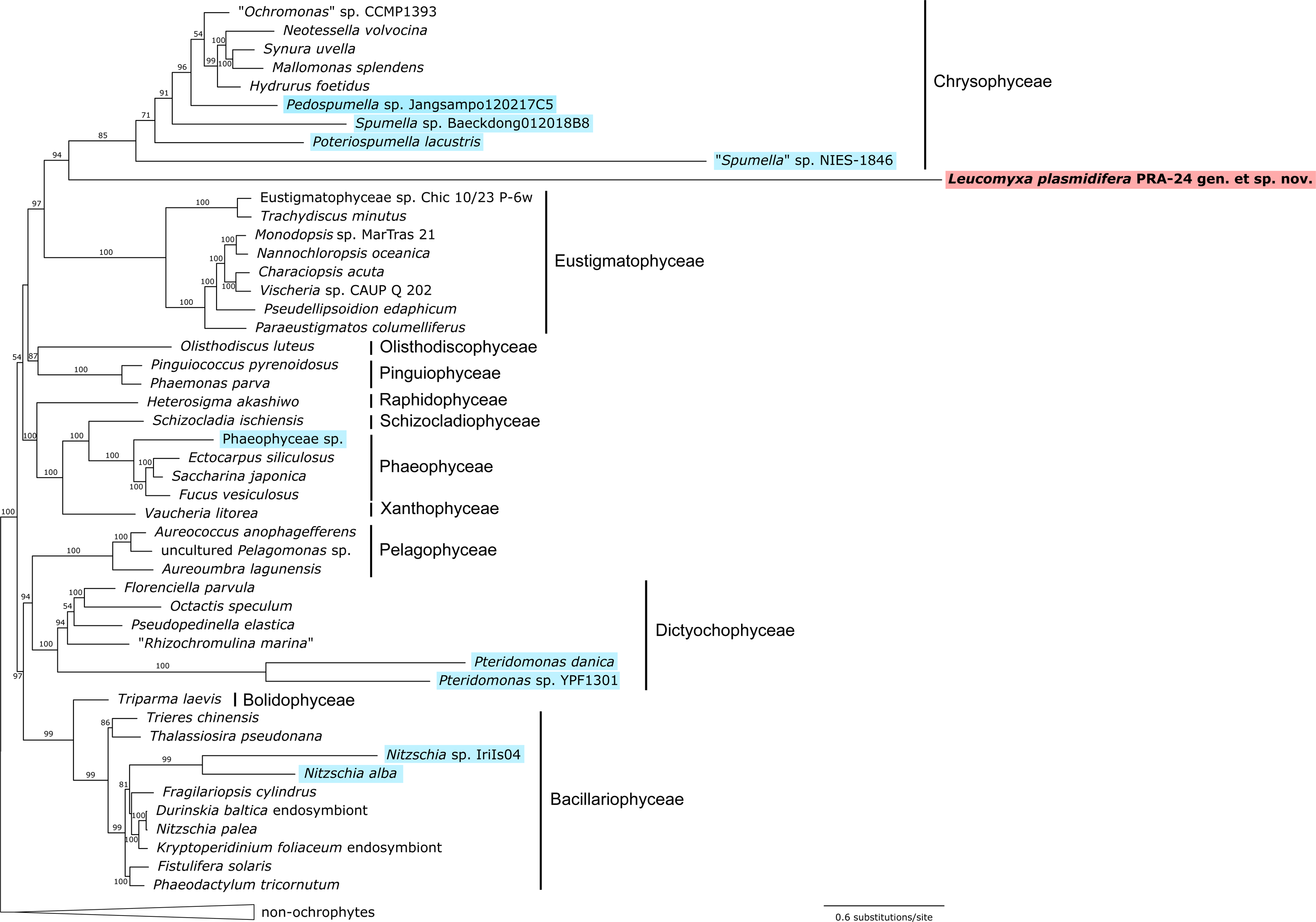
Phylogenetic relationships in the Ochrophyta based on plastid genome-encoded proteins. Displayed is the ML phylogenetic tree (IQ-TREE, substitution model LG+C60+F+G) inferred from a concatenated alignment of 30 conserved proteins (6,627 aligned amino acid position). Bootstrap support values are shown when ≥50. Non- photosynthetic taxa are highlighted with a coloured background. The sequence resources for the taxa included in the analysis are provided in supplementary electronic material, table S4.

While confirming we have identified a *bona fide* plastid genome of *Leuc. plasmidifera*, the tree additionally informs about a profound variation in the relative substitution rates in plastid genomes of different phylogenetic lineages. The long branches formed by all non-photosynthetic ochrophytes included in the tree (figure 4) indicate that the sequences of their plastid genes have been (on average) evolving much quicker than the homologous genes in their photosynthetic relatives. This is a common phenomenon observed in diverse non-photosynthetic taxa outside ochrophytes (e.g. Donaher et al. 2009; Záhonová et al. 2016; Preuss et al. 2020; Pánek et al. 2022), pointing to a general relaxation of evolutionary constraints on the remaining plastidial molecular components upon the loss of photosynthesis. The branch lengths of the different colourless ochrophytes in our tree differ substantially, presumably reflecting not only how much the evolutionary rate increased when photosynthesis was lost in the particular lineage, but also how recent the switch is. Notably, the most divergent among all ochrophyte plastid genomes sequenced is by far that one from *Leuc. plasmidifera* (figure 4), which might indicate that the loss of photosynthesis in its lineage is more ancient than analogous events in other colourless ochrophyte lineages. We note that the divergent nature of the plastid protein sequences of the non-photosynthetic taxa may be responsible for some problematic aspects of the tree obtained, including the incomplete statistical support for the monophyly of the class Chrysophyceae and of the sisterhood of chrysophytes and *Leuc. plasmidifera*, as well as for the apparently incorrect placement of the *Pteridomonas* lineage, which is expected to branch with *Pseudopedinella elastica* based on 18S rRNA phylogenies (Kayama et al. 2020), yet in our tree makes a sister lineage to all other dictyochophytes (figure 4). Future analyses with an improved taxon sampling and employment of ever more complex substitution models that can better account with the heterogeneity of the evolutionary process across lineages (see, e.g., Muñoz-Gómez et al. 2022), will provide more accurate results.

A closer look at individual genes in the *Leuc. plasmidifera* plastome revealed additional manifestations of rapid evolution. One example is the *rpoC2* gene, which encodes the β’’ subunit of the plastidial RNA polymerase. In *Leuc. plasmidifera* it is split into two ORFs in the same reading frame but separated by a termination codon. The split maps into an internal poorly conserved region and it is unlikely to sign pseudogenization, as the respective protein is essential for the function of RNA polymerase and no nucleus-encoded homolog that would potentially functionally compensate for this subunit was found in our *Leuc. plasmidifera* transcriptome assembly. Hence, we assume that the β’’ subunit is reconstituted in *Leuc. plasmidifera* from separately synthesized N- and C-terminal halves, and we annotate the respective gene parts *rpoC2N* and *rpoC2C*. It is notable that the *rpoB* gene, encoding the β’ subunit of the plastidial RNA polymerase, is similarly split in the plastome of the non-photosynthetic chrysophyte “*Spumella*” sp. NIES-1846 (Dorrell et al. 2019), pointing to a possible common evolutionary tendency to RNA polymerase subunit fragmentation in non-photosynthetic plastids in ochrophytes.

Another notable example of a peculiar plastidial gene in *Leuc. plasmidifera* is *clpC*, the only gene in the *Leuc. plasmidifera* plastome that is not directly related to gene expression. The encoded ClpC protein is a hexameric ATP-dependent chaperone involved in proteolysis in the plastid, functioning as part of a larger complex called the Clp protease system (Nishimura and van Wijk 2015). In its conventional structure the protein includes, from the N- to the C-terminus, the so-called N-domain followed by two AAA+ domains (Nishimura and van Wijk 2015). Variation to the organization of the plastidial *clpC* gene has, however, been noted before in two ochrophyte lineages, chrysophytes and eustigmatophytes (Ševčíková et al. 2015). These two groups share a *clpC* split whereby the region encoding the N-domain represents a separate gene (typically annotated *clpCN*); this feature has been interpreted as a synapomorphy supporting specific relationship of the two ochrophytes classes together constituting a clade called Limnista. In eustigmatophytes the *clpC* gene has been rearranged further by an additional split in the region between the two AAA domains yielding two separate genes (designated *clpC_A* and *clpC_B*). The *Leuc. plasmidifera* plastome contains a single region homologous to the *clpC* gene and thus annotated with this gene name, but the gene is short and the encoded protein in fact corresponds only to the C-terminal half of the conventional ClpC protein that contains the second AAA+ domain. No protein separately encoded by the *Leuc. plasmidifera* nuclear genome that would compensate for the missing parts of the conventional ClpC protein (N-domain and the first AAA+ domain) was identified by searches of our *Leuc. plasmidifera* transcriptome data, suggesting that a major modification in the functioning of the plastidial Clp protease system.

To test the notion that the elevated evolutionary rate observed for the plastid-coding genes in the *Leuc. plasmidifera* plastome holds also for non-coding genes, we carried out a phylogenetic analysis of the 16S rRNA gene. In addition to a set of sequences covering the whole diversity of ochrophyte lineages (plus appropriate non-ochrophyte taxa as an outgroup) that were extracted from previously sequenced plastid genomes or transcriptome assemblies, we included sequences of complete or partial 16S rRNA gene amplicons from PCR-based environmental surveys detected in the NCBI nr nucleotide sequence database and found to exhibit specific relationship to the 16S rRNA gene from *Leuc. plasmidifera* or *Synchroma pusillum*. The final phylogenetic tree (supplementary electronic material, figure S4) showed the *Leuc. plasmidifera* sequence to have the root-to-tip cumulative length substantially higher compared to the branches representing 16S rRNA sequences of other known non-photosynthetic ochrophytes included in the analysis. This finding demonstrates that the extreme sequence divergence is not restricted to protein-coding genes in the *Leuc. plasmidifera* plastome. Even more notable, however, is the fact that the *Leuc. plasmidifera* sequence was found to be part of a broader fully supported clade additionally comprised of environmental sequences with a comparable (in a few cases even higher) sequence divergence. This strongly indicates that they come from *Leuc. plasmidifera* relatives that are likewise non-photosynthetic (note that some of them were misidentified as sequences coming from uncultured bacteria by their authors). The whole *Leuc. plasmidifera*-containing non-photosynthetic clade was clustered together with sequences from non-photosynthetic diatoms and dictyochophytes instead of its expected position close to the sequence from *S. pusillum*, but this is an obvious result of the notorious long branch-attraction artefact affecting in phylogenies highly divergent sequences.

The three nearly identical 16S rRNA gene sequences making a tight cluster deeply diverged from yet specifically related the plastidial 16S rRNA from *Leuc. plasmidifera* (GenBank accession numbers LN565456.1–LN565458.1) represent eDNA clones that according to the metadata in the respective database records were obtained from a sample entitled “refuse dump Nest N12 layer 1” (investigated as part of an unpublished study of microbial communities associated with refuse dumps created by the leaf-cutter ant *A. colombica*).

Crucially, the same sample yielded two of the environmental 18S rRNA gene sequences (LN580971.1 and LN581250.1) belonging to the *Leucomyxa* clade (see figure 2 and supplementary electronic material, table S1). While the LN580971.1 sequence is relatively similar to the 18S rRNA gene sequences from *Leuc. plasmidifera*, the LN581250.1 sequence is more diverged, paralleling the relative position of the “LN565456.1–LN565458.1” group in the 16S rRNA gene tree. This correspondence indicates that the *Leuc. plasmidifera*-related eDNA sequences in both nuclear 18S and plastidial 16S rRNA gene trees define the same clade of non-photosynthetic ochrophytes, with *Leuc. plasmidifera* being its only presently known representative.

### 2.4. Transcriptome data corroborate the presence of a reduced plastid organelle in *Leucomyxa plasmidifera*

The plastid genome of the *Leuc. plasmidifera* plastid by itself did not provide any clue as to why the plastid has been retained by the organism: all the functionally annotated genes in the genome encode products have house-keeping roles required for the functioning of the plastid as such. Therefore, we set out to investigate the repertoire of plastid proteins encoded by the *Leuc. plasmidifera* nuclear genome. Since the genome assembly, from which we retrieved the plastid genome sequence, is highly fragmented with regard to the nuclear genome, we sequenced and assembled the transcriptome of the organism, and looked for encoded proteins that bear characteristics of targeting to a plastid. As an ochrophyte, *Leuc. plasmidifera* is expected to harbour a plastid with four bounding membranes, the outermost of which is continuous with the nuclear envelope and the ER (Dorrell and Bowler 2017). The route of nucleus-encoded proteins to the plastid in ochrophytes includes their co-translational transport into the ER, subsequent translocation via the second bounding membrane by the specialized ERAD-derived machinery called SELMA, and finally transport through the inner two membranes (equivalent to bounding membranes of the primary plastid) by an apparatus homologous to the conventional plastid protein import machinery consisting of the TOC and TIC complexes (Maier et al. 2015). Plastid-targeted proteins in ochrophytes thus possess a characteristic N-terminal presequence, called the bipartite targeting signals (BTS), consisting of a signal peptide (SP) to mediate translocation into the ER, followed by a transit peptide-like (TPL) region recognized by the SELMA, TOC, and TIC complexes (Patron and Waller 2007). Furthermore, as established by experimental studies on plastid protein targeting in diatoms, proteins destined to the stroma (as opposed to those to be retained in the periplastidial compartment) are expected to exhibit phenylalanine, tyrosine, tryptophan, or leucine residue right after the SP cleavage site (Gruber et al. 2007).

To ascertain that these expectations hold for *Leuc. plasmidifera*, we searched the transcriptome data for homologs of the key components of the SELMA, TOC, and TIC complexes, and indeed we found most of those looked for (supplementary electronic material, table S5). These included orthologs of both derlin-related proteins constituting the core of the SELMA complex (sDer1-1 and sDer1-2), an ortholog of the pore-forming protein central to the TOC complex (in ochrophytes called Omp85), and homologs of several components of the TIC complex (Tic22 and Tic110 identified with confidence). Like other ochrophytes, *Leuc. plasmidifera* also possesses a homolog of the protein PPP1 originally identified in the periplastidial compartment of the apicomplexan *Toxoplasma gondii* and shown to be critical for plastid protein import in this organism (Sheiner et al. 2011; Maier et al. 2015). While its precise function in ochrophytes remains unknown, PPP1 includes a region homologous to the C-terminal domain of a group of recently characterized red algal proteins implicated in plastid protein import as putative receptors recognizing the plastidial transit peptide (Baek et al. 2022), further strengthening the idea of PPP1 being a plastid protein-targeting factor in rhodophyte-derived secondary plastids in general, including *Leuc. plasmidifera*. Furthermore, *Leuc. plasmidifera* encodes an ortholog of the stromal processing peptidase that mediates removal of the N-terminal transit peptide like region to yield mature stromal proteins. These findings not only provide an independent evidence that *Leuc. plasmidifera* has a plastid organelle, but also justify the notion that plastid-targeted proteins in *Leuc. plasmidifera* will exhibit the characteristic BTSs at their N-termini/ To evaluate the features of the expected BTS of plastid-targeted proteins in *Leuc. plasmidifera* we searched its transcriptome assembly for homologs of conserved nucleus-encoded proteins involved in plastidial transcription and translation, which are processes certainly taking place in the stroma of the *Leuc. plasmidifera* plastid, owing to the presence of a plastid genome. Homologs of many components of these processes are readily identifiable and are unlike to function outside the plastid. With these assumptions in mind we collected 21 *Leuc. plasmidifera* proteins with unambiguously determined N-terminal sequences, excluding those expected to be dually targeted to the mitochondrion based on previous results, such as aminoacyl-tRNA synthetases (Gile et al. 2015). The set of reference *Leuc. plasmidifera* plastidial proteins included the sigma factor subunit of the plastidial RNA polymerase, supporting functionality of this enzyme despite the split of the plastome-encoded β’’ subunit (see above), translation factors, ribosomal proteins and factors implicated in plastid ribosome biogenesis (supplementary electronic material, table S5). All these proteins were predicted to contain a SP at the N-terminus, and as expected, the first amino acid residue following the SP cleavage site was usually phenylalanine, but also tyrosine, tryptophan, or leucine. However, in a few cases still another amino acid was predicted to be exposed at the N-terminus after SP cleavage, suggesting that the requirement for a F/Y/W/L residue is relaxed in *Leuc*. *plasmidifera* or that the SP cleavage sites are not always predicted correctly (indeed, the different prediction programs employed by us sometimes disagree with each other concerning the SP cleavage site; supplementary electronic material, table S5).

We thus conclude that plastid targeting of nucleus-encoded proteins in *Leuc. plasmidifera* generally follows the same rules as in other ochrophytes, and that bioinformatic identification of these proteins is possible with reasonable accuracy. Hence, we searched the *Leuc. plasmidifera* transcriptome assembly for homologs of components of common plastid-localised metabolic pathways and evaluated their likelihood of being targeted to the plastid by checking the nature of their N-termini. To consider the given *Leuc. plasmidifera* protein to be plastid-targeted, we required the presence of a signal peptide to be supported by at least three of the prediction programs (supplementary electronic material, table S5). However, we took some liberty in interpreting the outcomes of the prediction programs, considering as candidates for stromal proteins even some of those without a F/Y/W/L residue after the predicted SP cleavage if the protein is known to be stromal in other ochrophytes or plastid-bearing eukaryotes algae in general or if its presence in the stroma is expected based on the overall biochemical “logic” (for specific comments see supplementary electronic material, table S5). Given the general limitations of purely bioinformatic approaches, which seem to be particularly inept when it comes to recognizing dually localised proteins (Gould et al. 2023), the metabolic map of the *Leuc. plasmidifera* plastid presented below should be viewed as tentative.

In keeping with the non-photosynthetic and presumably highly reduced nature of the *Leuc. plasmidifera* plastid, the organism lacks enzymes of most of the metabolic pathways typical for a plastid or localised to it in other ochrophytes (Dorrell et al. 2017), including the type II fatty acid biosynthesis pathway, the DOXP (or MEP) pathway for the biosynthesis of isoprenoid precursors, and the pathways for biosynthesis of carotenoids, chlorophyll, plastoquinone, phylloquinone, and riboflavin. The Calvin-Benson-Bassham (CBB) cycle is also absent. In addition to the missing plastidial genes for RuBisCO subunits, no phosphoribulokinase homolog was identified in the transcriptome assembly. Homologs of the other CBB cycles do occur in *Leuc. plasmidifera*, but all seem to be cytosolic or mitochondrial based on the evaluation of the respective sequences by localisation predictions programs (supplementary electronic material, table S5). They thus obviously correspond to the enzymes of the glycolysis/gluconeogenesis (with some of the respective enzymes known to have mitochondrial isoforms in Stramenopiles; Río Bártulos et al. 2018) or the conventional cytosolic pentose phosphate pathway. The *Leuc. plasmidifera* also apparently lacks ferredoxin-NADP^+^ reductase and a plastid-targeted ferredoxin, the latter being a functionally versatile protein with a redox-active iron-sulfur (Fe-S) cluster and involved in multiple plastid-localised processes beyond photosynthesis itself (Hanke and Mulo 2013; Füssy et al. 2020). This is even more notable considering the fact that a ferredoxin-encoding gene (*petF*) has been retained by plastid genomes of several independently evolved non-photosynthetic chrysophytes (Dorrell et al. 2019; Kim et al. 2020).

Related to the lack of a plastidial ferredoxin is the absence in *Leuc. plasmidifera* of all components of the SUF system, a plastid-specific machinery for the assembly of Fe-S clusters. This is in a striking contrast with the presence of the SUF system in virtually all other plastids investigated, including those lacking any *suf* gene in their plastid genome (see also figure 3) and having the SUF system encoded fully by the nuclear genome. The latter concerns not only photosynthetic taxa (Przybyla-Toscano et al. 2021), but also eukaryotes bearing non-photosynthetic plastids, such as the parasitic trebouxiophyte *Helicosporidium* sp. (Pombert et al. 2014), various non-photosynthetic myzozoans (colpodellids, *Perkinsus*, *Platyproteum*; Janouškovec et al. 2015; Mathur et al. 2019), or *Euglena longa* (Novák Vanclová et al. 2020). Rare counterexamples exhibiting plastids devoid of the SUF pathway include the apicomplexan subgroup Piroplasmida (Sato 2011) and the ochrophyte *Pteridomonas* spp. (Kayama et al. 2020). Plastidial proteins, including those encoded by the nuclear genome, that require Fe-S clusters as their prosthetic groups acquire, to our knowledge, the clusters exclusively by the function of the SUF system inside the plastid.

Hence, the lack of the SUF system in *Leuc. plasmidifera* implies that the plastidial proteome in this organism is devoid of proteins depending of Fe-S clusters (a hypothesis to be tested with a future comprehensive reconstruction of the *Leuc. plasmidifera* plastidial proteome).

Plastid membranes generally contain four types of characteristic structural lipids, the phospholipid phosphatidylglycerol (PG), the galactolipids monogalactosyldiacylglycerol (MGDG) and digalactosyldiacylglycerol (DGDG), and the sulfolipid sulfoquinovosyldiacylglycerol (SQDG; Yoshihara and Kobayashi 2022). In plastid-bearing eukaryotes PG is synthesized by a plastid-localised pathway that is, however, typically parallel to a separate pathway producing this phospholipid for the use in other cellular compartments (Michaud et al. 2017). While *Leuc. plasmidifera* does have homologs of both enzymes dedicated to the PG biosynthesis (from the precursor CDP-diacylglycerol common to other phospholipids), none of them has a putative plastid-targeting presequence (supplementary electronic material, table S5). Completely absent from *Leuc. plasmidifera* are enzymes specific for the synthesis of MGDG and DGDG; indeed, the presence of galactolipids in non-photosynthetic plastids varies, with the functional significance of the differential presence of these lipids yet to be discerned (Botté and Maréchal 2014; Füssy et al. 2020). Somewhat unexpected is our identification in *Leuc. plasmidifera* of a homolog of the protein known as PHD1 and reported to be a putative plastid-specific UDP-glucose epimerase (UGE) catalysing interconversion of UDP-glucose and UDP-galactose, i.e. the latter being the source of galactose moieties in MGDG and DGDG (Li et al. 2011). Strikingly, the respective transcript sequence as represented in the transcriptome assembly is interrupted by two in-frame termination codons and differed at these positions from the corresponding gene sequence as represented in the genome assembly, but inspection of sequencing reads revealed that a substantial proportion of them mapping to the transcript support an alternative version differing by multiple single-nucleotide substitutions, including such that restore sense codons at the positions of both termination codons (supplementary electronic material, figure S5).

This could indicate the presence of two slightly different paralogs or allelic variants, with one being non-functionalized (a better genome assembly with a higher read coverage is needed to confirm this) and possibly indicating an on-going degradation of the corresponding metabolic process. We did not identify in *Leuc. plasmidifera* any other candidate for a plastidial enzyme that would use UDP-galactose as a substrate, leaving the physiological role of the UGE- catalysed reaction uncertain at present (although one possible explanation is discussed below).

In contrast to the apparent absence of galactolipid synthesis, the *Leuc. plasmidifera* plastid has evidently retained the synthesis of SQDG. Both critical enzymes, i.e. UDP-sulfoquinovose synthase (SQD1) and SQDG synthase (SQD2), are present in *Leuc. plasmidifera*, both bearing an N-terminal extension matching the structure of a BTP consistent with their presumed plastid localisation. The *Leuc. plasmidifera* plastid membranes are thus predicted to contain SQDG, similar to the leucoplasts of *E. longa*, non-photosynthetic diatoms, and some non-photosynthetic chrysophytes (Goddard-Borger and Williams 2017; Dorrell et al. 2019; Füssy et al. 2020). The synthesis of SQDG in the plastid relies on the provision of precursors. One is sulfite (SO3^2-^), in plastids commonly produced as part of the assimilatory sulfate reduction (Patron et al. 2008). Of the components of this pathway only a single candidate for a plastid-targeted enzyme was found in the *Leuc. plasmidifera* sequence data, phosphoadenosine phosphosulfate (PAPS) reductase, while the production of PAPS itself seems to be restricted to the cytosol in *Leuc. plasmidifera* (figure 5; supplementary electronic material, table S5). As we did not find any candidate for a sulfite reductase, which would catalyse the next step of sulfur assimilation towards cysteine, in the sequence data from the organism, the sole role of PAPS reductase in its plastid likely is provision of sulfite for the SQDG synthesis. The electrons for PAPS reduction are delivered by reduced thioredoxin, and indeed we identified a putative plastid-targeted variant of thioredoxin as well as of thioredoxin reductase in *Leuc. plasmidifera* (supplementary electronic material, table S5). The latter enzyme reduces thioredoxin in reaction with NADPH, and the latter electron carrier is also required for at least one additional reaction situated in the *Leuc. plasmidifera* plastid (the first step of the biosynthesis of haem catalysed by glutamyl-tRNA reductase, see the next section). However, our analyses did not reveal any plastid-localised NADPH source in *Leuc. plasmidifera*. How reducing equivalents are generated has not been elucidated even for the well-studied *Plasmodium* non-photosynthetic plastid (apicoplast), with import of NAD(P)H being considered as one of the hypothetical mechanisms (Niu et al. 2022).

**Figure 5.**
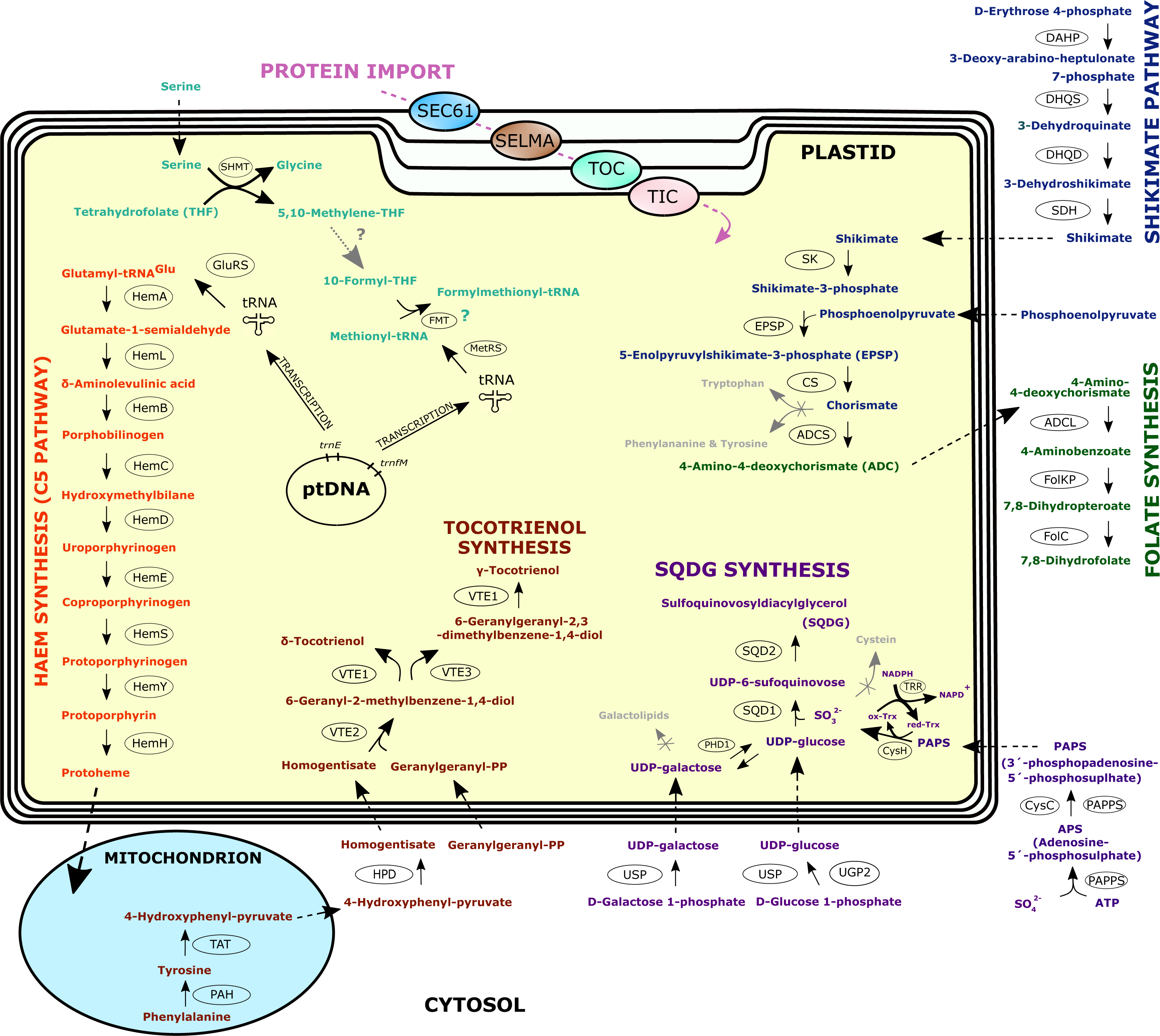
Bioinformatic reconstruction of the *Leucomyxa plasmidifera* plastid and its main metabolic pathways. The reconstruction is based on the data presented in supplementary electronic material, table S5. For simplicity all the putative plastid-localised reactions are shown such as to taking place in the plastid stroma, but it is likely that some of them (e.g., the haem synthesis pathway, see main text) occur in the periplastidial space (between the second and third plastid envelope membrane). The pathways are distinguished by different colours of the intermediates and final products. Arrows with solid lines indicate a chemical transformation (reaction), with the respective enzymes catalysing the reaction indicated in ovals adjacent to the arrows (for the meaning of the abbreviations used see below). The crossed arrows in grey (to Tryptophan, Phenylalanine & Tyrosine, Galactolipids, and Cysteine) highlights the apparent lack of these common plastidial biosynthetic branches in *Leuc. plasmidifera*, whereas the dotted arrow with a question mark leading from 5,10- Methylene-THF to 10-Formyl-THF indicates that the metabolic link is expected but could not be reconstructed by our analyses. The green question mark at the synthesis of Formylmethionyl-tRNA indicates that the process is expected to take place in the *Leuc. plasmidifera* plastid but not directly supported by results of localisation predictions for the respective enzyme (FMT). Arrows with dashed lines indicate transport of the particular compound between cellular compartments (for simplicity, the presumed import of diacylglycerol for SQDG biosynthesis from the ER is not illustrated). The complexes of the molecular machinery for the import of nuclear genome-encoded proteins into the plastid that have at least some of their subunits discernible in the transcriptome data from *Leuc. plasmidifera*, are illustrated in association with the respective membrane bounding the plastid (SEC61 = protein translocon at the ER; SELMA = translocon in the periplastidial membrane; TOC = translocon of the outer envelope membrane; TIC = translocon of the inner envelope membrane). Enzyme abbreviations: Haem synthesis – HemA = glutamyl-tRNA reductase; HemL = glutamate-1-semialdehyde 2,1-aminomutase; HemB = porphobilinogen synthase; HemC = hydroxymethylbilane synthase; HemD = uroporphyrinogen-III synthase; HemE = uroporphyrinogen decarboxylase; HemF = coproporphyrinogen III oxidase; HemY = protoporphyrinogen/coproporphyrinogen III oxidase; HemH = protoporphyrin/coproporphyrin ferrochelatase. Tocotrienol synthesis – PAH = phenylalanine- 4-hydroxylase; TAT = tyrosine aminotransferase; HPD = 4-hydroxyphenylpyruvate dioxygenase; VTE2 = homogentisate phytyltransferase / homogentisate geranylgeranyltransferase; VTE1 = tocopherol cyclase; VTE3 = MPBQ/MSBQ methytransferase. SQDG (sulphoquinovosyldiacylglycerol) synthesis – UGP2 = UTP--- glucose-1-phosphate uridylyltransferase; USP = UDP-sugar pyrophosphorylase; PHD1 = plastid-type UDP-glucose epimerase; PAPSS = 3’-phosphoadenosine 5’-phosphosulfate synthase; CysC = adenylylsulfate kinase; CysH = PAPS reductase; Trx = thioredoxin; TRR = thioredoxin reductase; SQD1 = UDP-sulfoquinovose synthase; SQD2 = sulfoquinovosyltransferase. Shikimate pathway – DAHP = 3-deoxy-D-arabino-heptulosonate 7-phosphate synthetase; DHQS = 3-dehydroquinate synthase; DHQD = 3-dehydroquinate dehydratase; SDH = shikimate dehydrogenase; SK = shikimate kinase); EPSP = 5- enolpyruvylshikimate-3-phosphate synthetase; CS = chorismate synthase. Folate synthesis – ADCS = 4-amino-4-deoxychorismate synthase; ADCL = 4-amino-4-deoxychorismate lyase; DHPS = dihydropteroate synthase (the activity conferred by the protein FolKP, which may in fact localise to the mitochondrion in *Leuc. plasmidifera*), DHFS = dihydrofolate synthase (the activity conferred by the protein FolC1). Folate-based one-carbon metabolism – SHMT = serine hydroxymethyltransferase; FMT = methionyl-tRNA formyltransferase. GluRS = glutamyl-tRNA synthetase; MetRS =methionyl-tRNA synthetase. Protein sequences corresponding to the enzymes and additional details are provided in see supplementary electronic material, table S5.

Another SQDG precursor is UDP-glucose, but we did not identify any candidate for a plastid-targeted enzyme that would mediate synthesis of this compound in *Leuc. plasmidifera*. This is understandable given the lack of an intraplastidial saccharide production in the *Leuc. plasmidifera* plastid and implies that the plastid imports UDP-glucose from the cytosol (putative cytosolic UDP-glucose-producing enzymes do exist in *Leuc. plasmidifera*; figure 5 and supplementary electronic material, table S5). However, a presently unknown mechanism of UDP-glucose synthesis may exist, as hypothesized for the plastid of the red alga *Cyanidioschyzon melorae* (Mori et al. 2018), and we cannot rule out the possibility that it also operates in the *Leuc. plasmidifera* plastid. An alternative explanation would rationalize the aforementioned presence of UGE in the *Leuc. plasmidifera* plastid, which could be involved in UDP-glucose generation from UDP-galactose if the latter can be imported in the *Leuc. plasmidifera* plastid. Finally, SQDG synthesis requires diacylglycerol, which is presumably delivered to the *Leuc. plasmidifera* plastid from the endoplasmic reticulum as part of the general exchange of lipids between these two compartments, as described from photosynthetic plastids (Block and Jouhet 2015).

Plastids, including the ochrophyte ones, are a site of production of a number of amino acids (Dorrell et al. 2017), but the *Leuc. plasmidifera* plastid seems to be depauperate in this regard (figure 5; supplementary electronic material, table S5). Thus, the organism seems to be incapable of *de novo* biosynthesis of aromatic amino acids (see also the next section), endowed only with a mitochondrion-targeted phenylalanine-4-hydroxylase that converts phenylalanine to tyrosine. *Leuc. plasmidifera* is also evidently auxotrophic for branched-chain amino acids (leucine, isoleucine, and valine). Of the enzymes of the pathway synthesizing lysine from aspartate, which in ochrophytes is wholly located in the plastid (Dorrell et al. 2017), only those catalysing the initial two and terminal two steps have homologs in *Leuc. plasmidifera*, with enzymes for the middle steps (five reactions) completely missing.

Furthermore, none of these *Leuc. plasmidifera* proteins has an N-terminal sequence that would be suggestive of its localisation in the plastid. Our analyses indicate that the first two enzymes serve in the synthesis of homoserine, a precursor of methionine (supplementary electronic material, table S5). The enzymes for the last two lysine biosynthesis steps, diaminopimelate epimerase (DapF) and diaminopimelate decarboxylase (DAPDC), seem to be true remnants of the ancestral ochrophyte plastid-localised pathway. The *Leuc. plasmidifera* DapF lacks any presequence and has thus been obviously relocated to the cytosol, whereas the prediction of DAPDC localisation is inconclusive, with the results of some programs compatible with it still being targeted to the plastid but others suggesting retargeting to the mitochondrion (supplementary electronic material, table S5). The function of DapF and DAPDC in *Leuc. plasmidifera* presumably is to allow utilization of food-derived lysine precursors (LL-2,6- and meso-2,6-diaminopimelate, both serving in bacteria also as peptidoglycan synthesis precursors) for the lysine production.

The only strong candidate for a plastidial enzyme directly involved in the generation of a particular amino acid we identified in *Leuc. plasmidifera* is serine hydroxymethyltransferase (SHMT), which catalyses conversion of serine to glycine with the extracted one-carbon group loaded onto tetrahydrofolate (or a backward reaction). Two other SHMT homologs, one mitochondrion-targeted and the other presumably cytosolic, exist in *Leuc. plasmidifera* (supplementary electronic material, table S5), so this reaction in the *Leuc. plasmidifera* plastid is expected to serve specifically the metabolism of the organelle. Like in other ochrophytes, serine itself is in *Leuc. plasmidifera* synthesized *de novo* (from 3-phosphoglycerate) outside the plastid, judging from the sequence features of the respective enzyme homologs. The physiological role of the plastidial SHMT thus seems to be feeding one-carbon units into the plastidial folate cycle using serine imported from the cytosol. A folate-based one-carbon cycle is expected to operate in the *Leuc. plasmidifera* as a source of formyltetrahydrofolate for the synthesis of formylmethionyl-tRNA (required for translation initiation in the plastid), but we could not identify any obvious candidates for plastid-localised enzymes that would mediate the conversion of methylenetetrahydrofolate to formyltetrahydrofolate, and even methionyl-tRNA formyltransferase (FMT) is present in *Leuc. plasmidifera* in only a single version, predicted to be mitochondrial (supplementary electronic material, table S5), where it is also needed for formylmethionyl-tRNA production. However, according to the respective record in the KEGG database (https://www.genome.jp/entry/2.1.2.9), a single FMT homolog is a rule rather than an exception among plastid-bearing eukaryotes, indicating that dual localisation of this enzyme to the mitochondrion and the plastid is common and likely occur also in *Leuc. plasmidifera*. No candidate was identified for a plastid-targeted cysteine synthase that would assimilate sulfide by its reaction with phosphoserine or O-acetyl-serine to make cysteine, which is consistent with the aforementioned absence of sulfite reductase in *Leuc. plasmidifera*. Concerning the two other amino acid biosynthesis enzymes operating in ochrophyte plastids, glutamate synthase appears to be completely absent from *Leuc. plasmidifera*, whereas only the mitochondrion-targeted isoform of glutamine synthetase is present in this organism.

### 2.5. Biosynthesis of haem, a folate precursor, and tocotrienols are the key physiological functions of the *Leucomyxa plasmidifera* plastid

All the processes discussed so far as localised to the *Leuc. plasmidifera* plastid serve directly in the biogenesis and maintenance of the organelle itself. This holds even for the SQDG synthesis, as no role of this structural lipid outside plastids has been defined in any eukaryote. So, what are then the key physiological functions of the plastid that underpin its retention by the *Leuc. plasmidifera* “host” cell? Our analyses of the transcriptome data indicate that the *Leuc. plasmidifera* plastid hosts of at least three metabolic pathway that clearly or potentially serve to the benefit of the whole cell (figure 5).

The first one is a pathway responsible for the synthesis of haem. All ochrophytes investigated so far possess a single haem biosynthesis pathway that is fully contained in the plastid and serves not only in the production of haem and other tetrapyrroles for the plastid itself, but supplies with haem the cell as a whole (Cihlář et al. 2019). It was, therefore, not surprising to find in the *Leuc. plasmidifera* transcriptome assembly homologs of all enzymes of the pathway from glutamyl-tRNA reductase (HemA) and glutamate-1-semialdehyde 2,1- aminomutase (HemL) to the final protoporphyrin ferrochelatase (HemH). The presence of

HemA and HemL implies that the key intermediate of the haem biosynthesis, 5- aminolevulinate, is created in *Leuc. plasmidifera* via the so called C5 pathway characteristic for plastids and using glutamyl-tRNA as the haem precursor (Cihlář et al. 2019). This means that Glu-tRNA specified by the *Leuc. plasmidifera* plastid genome serves not only in translation in the plastid, but also is critical for haem production. The alternative route to 5- aminolevulinate, the so-called C4 (or Shemin) pathway based on the reaction of glycine and succinyl-CoA catalysed by the enzyme 5-aminolevulinate synthase (ALAS) localised to the mitochondrion in eukaryotes is absent from *Leuc. plasmidifera*, as we could not detect any ALAS homolog in its transcriptome assembly. This is consistent with the general absence of this enzyme from ochrophytes (Cihlář et al. 2019). As expected, all the HemA to HemH enzymes carry a plastid-targeting presequence, but strikingly, the prediction programs are virtually fully consistent in predicting the SP cleavage sites such that they are not followed by any of the F/Y/W/L amino acid residue (supplementary electronic material, table S5). This may be an indication that rather than to the plastid stroma, the haem biosynthesis pathway in *Leuc. plasmidifera* localises to the periplastidial space. This would, however, pose a challenge to the function of at least the first enzyme of the pathway, HemA, which needs an access to its substrate glutamyl-tRNA certainly generated in the plastid stroma. Hence, we leave the exact special partitioning of the haem biosynthesis pathway in the *Leuc. plasmidifera* plastid open. Evidence for possible ramifications of the haem biosynthesis pathway that would localise to the *Leuc. plasmidifera* plastid, such as the branches leading to sirohaem or chlorophylls, or further modifications of heam towards bilins, was not identified in our data. While possible function of haem in the *Leuc. plasmidifera* plastid cannot be excluded at the present stage of knowledge, we infer that the primary clients of the plastid-localised haem synthesis are cytochromes of the mitochondrional respiratory chain.

Plastids commonly house the shikimate pathway supplying precursors of aromatic compounds, including aromatic amino acids and folate (Richards et al. 2006; Tzin et al. 2012). Our analysis of the *Leuc. plasmidifera* transcriptome assembly revealed that this organism has homologs of all seven enzymes of the pathway (supplementary electronic material, table S5). The protein corresponding to the first enzyme of the pathway, 3-deoxy-D-arabino-heptulosonate 7-phosphate synthetase catalysing condensation of phosphoenolpyruvate and D-erythrose 4-phosphate, does have an N-terminal extension compared to the targeting signal-free prokaryotic homologs, but a possible signal peptide was not recognized by any of the prediction programs employed and possible mitochondrial targeting presequence was predicted inconsistently (only by some of the programs and with weak scores only; supplementary electronic material, table S5), so we interpret the enzyme to most likely be cytosolic (figure 5). The protein representing the second enzyme of the pathway, 3-dehydroquinate synthase, has no presequence and the prediction programs do not suggest any specific targeting (supplementary electronic material, table S5), consistent with the notion that this enzyme is cytosolic in *Leuc. plasmidifera*.

The following two enzymes of the shikimate pathway, 3-dehydroquinate dehydratase and shikimate dehydrogenase, are parts of a single fusion protein possessing an N-terminal extension compared to bacterial homologs, but it is not recognised as any characteristic targeting presequence by any of the programs used by us, again suggesting cytosolic localisation of the protein. In contrast, the homologs of the remaining three enzymes, namely shikimate kinase, 5-enolpyruvylshikimate-3-phosphate synthetase (also called 3- phosphoshikimate 1-carboxyvinyltransferase), and chorismate synthase, are all predicted to be targeted to the plastid. Thus, the organisation of the shikimate pathway in *Leuc. plasmidifera* (figure 5) seems to differ from that in other ochrophytes, including the non-photosynthetic members of the diatom genus *Nitzschia*, which contain the whole pathway in the plastid (Kamikawa et al. 2017). The relocation of the initial steps of the pathway from the plastid in *Leuc. plasmidifera* may relate to the loss of the Calvin-Benson cycle and hence a source of D-erythrose 4-phosphate in the organelle, which instead is apparently provided by the cytosolic pentose phosphate pathway in *Leuc. plasmidifera*.

The final product of the shikimate pathway, chorismate, is a branching point leading to multiple aromatic compounds. One branch, starting with the enzyme chorismate mutase, leads to phenylalanine and tyrosine. However, the *Leuc. plasmidifera* transcriptome assembly lacks any discernible homologs of this or other enzymes of phenylalanine and tyrosine biosynthesis. Another branch leads to tryptophan, with the first two reactions being catalysed by anthranilate synthase and anthranilate phosphoribosyltransferase. We did not detect any candidates for these enzymes in the *Leuc. plasmidifera* transcriptome data, so we were surprised to find a protein comprising regions of homology to the two following enzymes of the tryptophan biosynthesis pathway, i.e. phosphoribosylanthranilate isomerase (TrpF) and indole-3-glycerol phosphate synthase (TrpC), fused in the reversed order from the N- to the C-terminus (supplementary electronic material, table S5). However, a closer inspection of the amino acid sequence of both the TrpF and TrpC region revealed mutations at invariant positions that were identified as critical for the catalytic activity of both enzymes: a cysteine residue, crucial for the TrpF function (Henn-Sax et al. 2002) is mutated to a serine reside in the *Leuc. plasmidifera* protein (supplementary electronic material, figure S6A), whereas a lysine residue essential for the catalytic activity of TrpC (Darimont et al. 1998) is mutated to a alanine residue in the *Leuc. plasmidifera* protein (supplementary electronic material, figure S6B). These changes suggest that the *Leuc. plasmidifera* TrpC-TrpF fusion protein is catalytically inactive, i.e. that it is a pseudoenzyme (Jeffery 2019) with presently undefined function in the *Leuc. plasmidifera* plastid. Since *Leuc. plasmidifera* apparently lacks candidates for the remaining enzyme of tryptophan biosynthesis, we conclude that this organism is auxotrophic for trypthophan as well as phenylalanine and tyrosine.

The only metabolic sink for chorismate produced in the *Leuc. plasmidifera* plastid thus seems to the biosynthetic branch leading to folate. Specifically, we identified in the *Leuc. plasmidifera* transcriptome assembly homologs of all enzymes of the pathway, with the one catalysing the first step, 4-amino-4-deoxychorismate synthase, having a putative plastid-targeting presequence (supplementary electronic material, table S5). Enzymes for the following steps, from 4-amino-4-deoxychorismate lyase to the bifunctional dihydrofolate synthase / folylpolyglutamate synthase, are most likely all localised outside the plastid based on the features of their N-termini. Hence, the *Leuc. plasmidifera* plastid seems to a site of production of 4-amino-4-deoxychorismate, which is then exported to the cytosol for further processing up to the one-carbon carrier dihydrofolate (figure 5).

The third pathway, whose presence in the *Leuc. plasmidifera* plastid can be reconstructed based on the transcriptomic data, may look surprising for a non-photosynthetic plastid, as it is commonly associated with photosynthesis. Namely, *Leuc. plasmidifera* possesses homologs of three enzymes of the tocochromanol (i.e. tocopherol and tocotrienol) biosynthesis pathway (Mène-Saffrané 2017), all with predicted plastidial localisation: homogentisate phytyltransferase / homogentisate geranylgeranyltransferase, MPBQ/MSBQ methyltransferase, and tocopherol cyclase (supplementary electronic material, table S5). The first enzyme catalyses the first committed step of the pathway, prenylation of homogentisate with a phytyl or geranylgeranyl residue (using phytyl-PP or geranylgeranyl-PP as the source), initiating thus synthesis of tocopherols or tocotrienols, respectively. *Leuc. plasmidifera* possesses both enzymes required for the production of homogentisate from L-tyrosine, predicted to be localised in the mitochondrion and the cytosol, respectively, implying homogentisate import into the plastid (figure 5; supplementary electronic material, table S5). Geranylgeranyl-PP is a ubiquitous metabolite and *Leuc. plasmidifera* contains a single form of the enzyme catalysing its production (geranylgeranyl diphosphate synthase, type III), which is apparently cytosolic (supplementary electronic material, table S5). Phytyl-PP is a specific derivative of geranylgeranyl-PP synthesised by the action of geranylgeranyl diphosphate reductase (GGDR), but no candidate for this enzyme is discernible in the *Leuc. plasmidifera* sequence data. This organism thus probably lacks the capability of synthesizing phytyl-PP, which is consistent with the lack of other pathways utilising this compound (chlorophyll and phylloquinone biosynthesis).

Based on these findings we infer that *Leuc. plasmidifera* produces tocotrienols but not tocopherols, specifically γ- and δ-tocotrienol, which assumes import of geranylgeranyl-PP from the cytosol (figure 5). It should, however, be noted that *Euglena* spp., while containing phytyl chains in chlorophyll and phylloquinone, also lack a discernible GGDR homolog (Novák Vanclová et al. 2020; Füssy et al. 2020), suggesting the existence of an alternative, currently unknown form of the enzyme that may theoretically be present also in *Leuc. plasmidifera*. The enzyme for the additional methylation step converting γ- and δ-tocotrienol (or γ-/δ-tocopherol) to α- and β-tocotrienol (or α-/β-tocopherol), i.e. tocopherol O-methyltransferase, seems to be absent from *Leuc. plasmidifera* (we could not retrieve it even when using as the query the tocopherol O-methyltransferase sequence from the closely related photosynthetic species *Synchroma pusillum*). The physiological role of γ- and δ-tocotrienols in a non-photosynthetic plastid is unclear, but they may be used in other parts of the cell – especially mitochondria and peroxisomes as general lipophilic antioxidants protecting membrane lipids against reactive oxygen species. Interestingly, a recent study of the non-photosynthetic euglenophyte *Euglena longa* provided bioinformatic as well as direct biochemical evidence for the production of tocopherols in this organism (Füssy et al. 2020), so the situation we have encountered in *Leuc. plasmidifera* is not unprecedented.

### 2.6. The *Leucomyxa plasmidifera* mitochondrial genome includes an unusual long insertion in the *cob* gene

In addition to the plastid genome, the genomic data we gathered from the *Leuc. plasmidifera* culture yielded a complete mitochondrial genome of the organism, recovered as a single gapless scaffold with sequence identity of the 5’ and 3’ termini consistent with the expected circular-mapping structure of the genome. The *Leuc. plasmidifera* mitogenome is 44,072 bp in length and contains 48 putative protein-coding genes, including 12 ORFs that do not correspond to standard mitochondrial genes or are too divergent to be identified as such, in addition to 26 tRNA genes and three rRNA genes (figure 6A; supplementary electronic material, table S6). The latter gene category includes a gene for 5S rRNA that is frequently not found or annotated in mitochondrial genomes (Valach et al. 2014). The set of tRNAs specified by the mitogenome is as typical for ochrophytes and stramenopiles as a whole, including all expected species needed to translate the codons for all amino acids except for threonine (the respective tRNA gene was most likely lost already in the stramenopile stem lineage and is presumably compensated for by importing the missing tRNA from the cytosol; Ševčíková et al. 2016; Sibbald et al. 2021). Consistent with the nature of the mitochondrial tRNA gene set, no departures from the standard codon meaning in the *Leuc. plasmidifera* mitochondrion were indicated by a FACIL analysis (supplementary electronic material, figure S3B).

**Figure 6.**
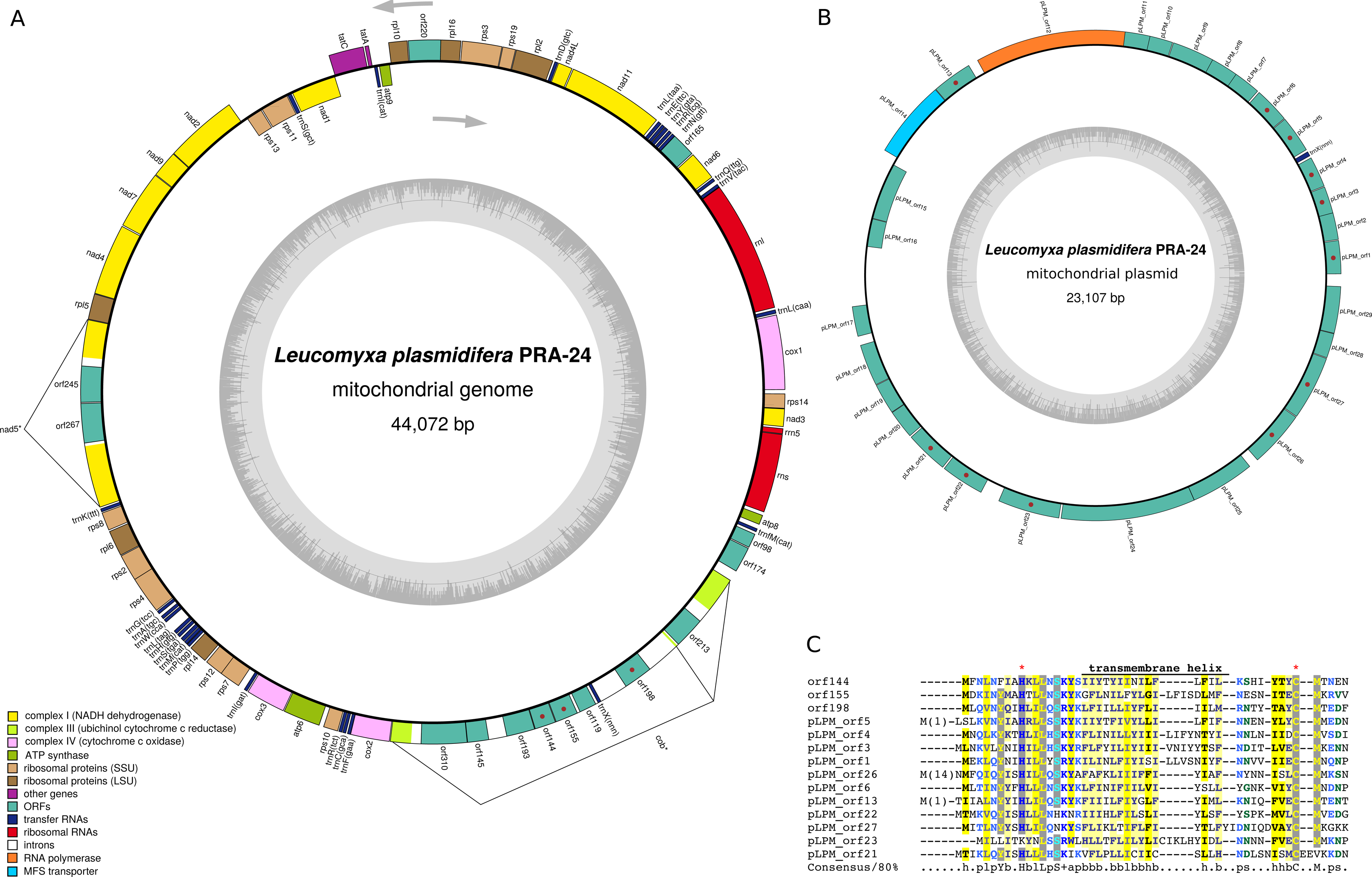
Mitochondrial genome and plasmid of *Leucomyxa plasmidifera*. (A) A schematic map of the mitogenome. The display convention is the same as for the plastid genome map (figure 3A). Note the *nad5* and *cob* genes interrupted with one and two ORF-containing introns, respectively, with the first intron in the latter gene being very long and containing even a tRNA gene (see the text for further details). (B) A schematic map of the mitochondrial plasmid (pLPM). (C) A novel conserved domain shared by multiple proteins encoded by pLPM and the putative plasmid-derived insert in the mitochondrial *cob* gene intron. ORFs encoding the proteins with this domain are highlighted with a red dot in the maps in parts A and B.

We identified in the *Leuc. plasmidifera* mitogenome most of the protein-coding genes known to occur in previously characterized ochrophyte mitogenomes (supplementary electronic material, table S6). In a few cases the identification required the highly sensitive profile HMM-HMM comparisons by HHpred (*rpl2*, *rpl5*), and in one case (*rpl10*) the identification is tentative, based on the similarity of a *de novo* structural model of the encoded protein obtained with AlphaFold2 to the ribosomal L10 protein (as assessed by FoldSeek). It is possible that some of the few remaining ribosomal protein genes occurring in at least some of the other ochrophyte mitogenomes yet not found in *Leuc. plasmidifera*, namely *rpl31* and *rps1*, are in fact present but have diverged beyond recognition with the currently available computational tools, as the mitogenome includes three unidentified ORFs (*orf98*, *orf174*, and *orf220*; putting aside intronic ORFs, see below). On the other hand, the *Leuc. plasmidifera* mitogenome certainly lacks the *atp1* gene, which in a previous study was hypothesized to have been preserved only in eustigmatophytes among all ochrophyte groups. Indeed, the *Leuc. plasmidifera* transcriptome records the existence of a nucleus-encoded mitochondrion targeted version of the Atp1 protein (supplementary electronic material, table S5), as is common for all non-eustigmatophyte ochrophytes investigated so far (Ševčíková et al. 2016).

A notable feature of the *Leuc. plasmidifera* mitogenome is the presence of three introns, one in *nad5* (group IB intron) and two in *cob* (both group ID introns). No introns have so far been reported in these two genes from stramenopile mitochondria, and we did not detect any other case when searching stramenopile mitochondrial genomes available in GenBank, indicating gain of these introns in the *Leuc. plasmidifera* lineage. The *nad5* intron and the second *cob* intron in *Leuc. plasmidifera* each encode a LAGLIDADG family homing endonuclease (LHE), i.e. a protein implicated in intron mobility by intron homing (Mukhopadhyay and Hausner 2021). BLASTP searches indicate the highest similarity of both *Leuc. plasmidifera* mitochondrial LHEs to proteins encoded by fungal mitochondria, suggesting that the respective introns including the LHE-encoding ORFs were gained by horizontal gene transfer (HGT) presumably from fungal sources.

While the LHE-encoding *nad5* and *cob* introns are more or less conventional, the first *cob* intron has a surprising form: it is exceptionally long (5,799 bp) and contains not only multiple ORFs (seven, when ORFs with ≥100 codons are considered), but also a tRNA gene. To ascertain that this long *cob*-interrupting region behaves as a *bona fide* intron, we mapped available RNAseq reads onto the mitochondrial genome sequence and indeed found multiple reads supporting the existence of a contiguous RNA molecule with the long intron removed. While the overall arrangement of the *cob* gene would be compatible with the first intron spliced out by the conventional *cis*-splicing mechanism, its sheer length raises the question whether an alternative interpretation should be invoked. Specifically, it is conceivable that the full-length *cob* mRNA is created from two separately transcribed regions (one including the first exon and the other including the remaining two exons) by the process of *trans*-splicing. Indeed, group I intron *trans*-splicing has been described from mitochondria of various eukaryotes (Mukhopadhyay and Hausner 2021).

None of the seven protein sequences deduced from the ORFs in the first *cob* intron retrieved any significant hit in BLASTP searches against major repositories of protein sequences (NCBI nr protein sequence database, EukProt) or when analysed with HHpred. TBLASTN searches against the transcriptome assembly from *Synchroma pusillum*, the closest *Leuc. plasmidifera* relative with genome-scale sequence data available, retrieved no significant hits either. It is, however, notable that six of the seven mysterious proteins exhibit predicted transmembrane helices: one (Orf144, Orf155, and Orf198; see also the next section), two (Orf145), and five (Orf310). The tRNA gene in the intron is annotated by MFannot as *trnL(uaa)*, but another *trnL(uaa)* gene occurs elsewhere in the genome and that one apparently corresponds to the standard vertically inherited mitochondrial gene as inferred from its high similarity to *trnL(uaa)* genes in other ochrophyte mitogenomes. The identity of the standard *trnL(uaa)* gene is additionally confirmed by it matching with the highest score (73.4) the Leu-tRNA isotype in a tRNAscan-SE search against bacterial tRNA models. In contrast, the intronic *trnL(uaa)* matches with the highest score (47.2) the Thr-tRNA isotype, whereas the match to the Leu-tRNA isotype is very weak (the score of 18.9). Furthermore, the standard *trnL(uaa)* exhibits a variable arm, a characteristic expected for a Leu-tRNA (Giegé and Eriani 2023), whereas the predicted secondary structure of the intron-specified tRNA lacks it (supplementary electronic material, figure S7A, B). Hence, the identity and functionality of the latter tRNA is uncertain and we annotate the genes as *trnX(uaa)*.

### 2.7. A putative novel mitochondrial circular plasmid occurs in *Leucomyxa plasmidifera*

Although the exact nature and function of the long intronic region in the *cob* gene remains unknown, we obtain a hint concerning its origin. While no homologs of the seven ORFs contained in the intron could be identified in public databases (see above), four of them did give significant non-self hits (E-value >1e-5) when used as queries in TBLASTN searches against the whole genome assembly of *Leuc. plasmidifera*. Interestingly, all these hits corresponded to the same scaffold of 23,184 bp with relatively very high read coverage (194×, compared to 58.4× for the mitogenome, 13.2× for the plastome, and only ∼2× for the nuclear genome) and with a region (77 bp) at the very 5’ end identical to a region at the very 3’ end. These features indicate that the scaffold corresponds to a multicopy circular-mapping DNA molecule of the total size of 23,107 bp. When this scaffold was compared with BLASTN against the whole *Leuc. plasmidifera* genome assembly, a single non-self significant hit (E-value of 4e-21) was found, represented by a 74 bp-long match (93% identity) to the *Leuc. plasmidifera* mitogenome scaffold. The match mapped to a putatively non-coding region within the long first intron in the *cob* gene downstream of the intronic tRNA gene, so it represented another link between the mitogenome and the high-coverage unknown genetic element.

These observations suggested that we encountered a putative circular DNA molecule occurring in the *Leuc. plasmidifera* mitochondrion, distinct from the mitogenome itself, and related to or descended from a DNA source of the insertion that have extended the first *cob* intron. As such, this genetic element is reminiscent of some previously described mitochondrial plasmids, which have been reported to have the ability to integrate to the mitochondrial genome or to exhibit regions of high sequence similarity shared with mitochondrial genomes of the same organism (Hausner 2012; Swart et al. 2012; Nishimura et al. 2019). Mitochondrial plasmids have been identified in fungi, plants, the amoebozoan *Physarum polycephalum*, ciliates, and a centrohelid, and are usually linear, but circular forms are known as well. Therefore, we interpret the multicopy DNA element found in *Leuc. plasmidifera* as a novel mitochondrial circular plasmid, the first to be reported from a stramenopile. We call the plasmid pLPM (with the abbreviation reflecting the species name and the putative mitochondrial localisation). The immediately apparent salient feature of pLPM is its size far exceeding the largest mitochondrial plasmid we could identify in the GenBank database: the linear plasmid pTB_1 from the fungus *Termitomyces bulborhizus* reaching the length of 15,455 bp including terminal inverted repeats (MW874159.1). Among the previously reported circular mitochondrial plasmids the largest seem to be pVS from *Neurospora intermedia* isolate M1991-107 (AY553873.1) with the length of 3,774 bp, further underscoring the novelty of pLPM.

Considering ORFs with the minimal length of 300 bp (=100 amino acid residues), there are 29 of them in the plasmid, generally densely packed (sometimes even with short overlaps) and except two all in the same direction (figure 6B). The plasmid additionally contains a single tRNA gene, a non-coding segment of 74 bp matching with a few substitutions a non-coding region in the long *cob* intron in the mitogenome (see above), and a few regions with no specific annotation, some of which contain short potentially functional but non-conserved ORFs that we omitted from the annotation. Of the proteins encoded by the annotated ORFs only one (*pLPM_orf14*) retrieved a significant hit (E-value <1e-5) when its putative protein product was compared to the NCBI *nr* database with BLASTP. Strikingly, this protein is a member of the major facilitator superfamily (MFS) transporters, with the highest similarity hits (amino acid identity up to 40%) coming predominantly from (putatively) endosymbiotic bacteria (metagenomically assembled genomes assigned to Rickettsiaceae or various member of Holosporales, such *Candidatus* Paracaedimonas acanthamoebae). It is thus likely that the MFS transporter encoded by *pLPM_orf14* was acquired by the plasmid via horizontal gene transfer from a bacterial endosymbiont co-occurring in the same host. Considering the general characteristics of MFS proteins (Drew et al. 2021), it is expected that the *pLPM_orf14* product mediates transport of specific metabolites into or from the *Leuc. plasmidifera* mitochondrion, although predicting its substrate specificity bioinformatically is beyond current possibilities. At any rate, the existence of such a gene in pLPM is noteworthy, as to our knowledge there is no previous report for a mitochondrial plasmid encoding a protein with a metabolic function.

Using HHpred searches as a more sensitive homology detection approach, only one additional protein encoded by ORFs in pLPM could be assigned to a previously defined gene family, namely the putative *pLPM_orf12* product (with its 769 amino acid residues the longest pLPM-encoded protein), which in HHpred searches gave a highly significant hit (E-value <1e-99) to a group of proteins typified by T3/T7 (T-odd) phage RNA polymerases. This group additionally includes the nucleus-encoded mitochondrial RNA polymerase (POLRMT) responsible for transcription of mitochondrial genes as well as, notably, RNA polymerases encoded by diverse previously identified mitochondrial plasmids (Shutt and Gray 2006; Hausner 2012; Nishimura et al. 2019). While simple homology searches were sufficient to identify many mitochondrial plasmid-encoded RNA polymerases due to their sequence conservation, the highly divergent nature of the *Leuc. plasmidifera* sequence is not unprecedented, as documented by a similar case of the plasmid-encoded RNA polymerase in the ciliate *Oxytricha trifallax* (Swart et al. 2012). The *Leuc. plasmidifera* transcriptome assembly contains a transcript encoding a close homolog of POLRMT from other eukaryotes possessing a predicted N-terminal mitochondrial transit peptide (supplementary electronic material, table S5), which is thus the obvious *bona fide* mitochondrial RNA polymerase mediating transcription of the *Leuc. plasmidifera* mitogenome. The divergent RNA polymerase homolog encoded by pLPM is then presumably dedicated to transcribing genes harboured by the plasmid itself.

Further analyses of the ORFs contained in pLPM and the long intron in the mitochondrial *cob* gene revealed that eleven of the former and three of the latter encode proteins sharing a homologous region. As apparent from a multiple sequence alignment of all 14 sequences, the proteins contain a conserved short N-terminal domain (∼40 amino acid residues) followed by a prolonged non-conserved C-terminal extension (figure 6C; supplementary electronic material, figure S8). The central segment of the N-terminal domain is predicted to form a transmembrane helix. The region upstream of the helix contains a nearly absolutely conserved histidine residue (with one exception), whereas the region downstream contains an absolutely conserved cysteine residue (nearly always as part of a motif “CM”). A profile HMM derived from the alignment of the 14 available sequences representing the domain did not reveal any homologs outside *Leuc. plasmidifera* when used as a query in HHMER searches against the NCBI nr protein sequence database and the EukProt v3 database. Using the same profile HMM as a HHpred query to perform a more sensitive homology search based on an HMM-HMM comparison did not retrieve any significant hit either. The function of this novel domain and the whole protein family encoded by the *Leuc. plasmidifera* mitogenome and pLPM thus remains truly enigmatic.

One additional ORF from the long *cob* intron, *orf119*, proved to share homology with a region of pLPM, namely the one corresponding to *pLPM_orf28* (pairwise BLASTP comparison of the encoded proteins showing the E-value of 9e-27). A profile HMM derived from the alignment of the two homologous proteins was used for HMMER and HHpred searches to identify potential additional homologs in *Leuc. plasmidifera* itself or other organisms, but without retrieving any credible hit. Two additional pairs of homologs (pairwise BLASTP comparison with the E-values of 7e-11 and 4e-07, respectively) were identified among the proteins encoded by the plasmid alone, specifically the gene pairs *pLPM_orf7*/*pLPM_orf19* and *pLPM_orf8*/*pLPM_orf20*, but again no additional homologs in *Leuc. plasmidifera* or elsewhere could be detected by searches employing profile HMMs derived from the alignments of the respective pairs of proteins. The remaining 13 proteins putatively encoded by the plasmid remain true orphans with no homolog discerned by any method employed, but it is notable that two of them *pLPM_orf25* and *pLPM_orf24*, are predicted to contain one and two transmembrane helices, respectively. There is presently no indication that pLPM would encode a DNA polymerase, a protein frequently encoded by mitochondrial plasmids found in other taxa and presumably involved in their propagation (Hausner 2012; Swart et al. 2012; Nishimura et al. 2019). It is thus likely that pLPM replication depends on the nucleus-encoded DNA polymerase that primarily mediates replication of the mitochondrial genome in stramenopiles, i.e. POP (Harada et al. 2024; supplementary electronic material, table S5).

The single tRNA specified by pLPM is unusual, reminiscent of the tRNA specified by the long *cob* intron in the *Leuc. plasmidifera* mitogenome, although the two tRNAs are not particularly similar to each other. The annotation and predicted secondary structure of the plasmid-specified tRNA provided by tRNAscan-SE depends on the tRNA model set used for the analysis (supplementary electronic material, figure S7C). With the “other mitochondrial” set the tRNA is predicted with an undefined anticodon sequence and the anticodon loop being expanded by one nucleotide compared to the standard tRNA structure, i.e. being 8 rather than 7 nucleotides long. With the “bacterial” set the tRNA is predicted as being of the Ile-tRNA type with the anticodon UAU yet with the anticodon stem being interrupted by an insertion of three unpaired nucleotides. Furthermore, the acceptor stem has only six rather than canonical seven paired nucleotides, lacking the top pair. It is noteworthy that tRNA genes were previously identified in some fungal and plant mitochondrial plasmids, and that in some cases the gene for the respective tRNA species has been lost from the mitogenome of the same organism, indicating dependence of the mitochondrial translation on the plasmid-specified tRNA (Nieuwenhuis et al. 2023). The only essential tRNA species lacking a corresponding gene in the *Leuc. plasmidifera* mitogenome is the one needed to decode threonine codons, like in other stramenopiles presumably imported from the cytosol (see above). The plasmid-specified tRNA (or, for that matter, the tRNA specified by the pLPM-related *cob* intron) does not seem to be cognate to threonine codons (ACN), so its functionality remain uncertain.

## 3. **Conclusions and perspectives**

Before we started the work reported in this paper, *Leukarachnion* sp. PRA-24 was a candidate for an ochrophyte that has secondarily lost the plastid organelle. Confirming this would open up an exciting and rare opportunity to study the process of complete organelle elimination from a eukaryotic cell, but our results clearly demonstrate that this organism, here formally described as the new genus and species *Leucomyxa plasmidifera*, is more conventional in this regard and joins a growing list of secondarily non-photosynthetic eukaryote lineages that have retained a residual plastid. This conviction is based on the very strong genomic signature for the presence of a plastid organelle in *Leuc. plasmidifera*, including the plastid genome and numerous nuclear genes that encode hallmark plastid proteins, while unquestionable cytological evidence for the plastid is yet to be provided. Our identification of environmental sequences related to the *Leuc. plasmidifera* plastidial 16S rRNA gene and exhibiting a similarly extreme divergence indicate the existence of a broader ochrophyte clade characterized by a genome-carrying non-photosynthetic plastid. Organisms of this clade seem to be common in organic-rich terrestrial habitats and may represent an overlooked important component of soil microbiota that needs to be investigated further by targeted sampling and culturing. It is also important to settle the classification of *Leuc. plasmidifera* and the whole clade in Ochrophyta, particularly to resolve its systematic position with respect to the previously established class Synchromophyceae. A phylogenomic analysis utilizing the *Leuc. plasmidifera* transcriptome data generated here, combined with new data from organisms belonging to the phylogenetic neighbourhood of *Leuc. plasmidifera*, is underway and should provide a basis for a refined classification of the whole major ochrophyte branch.

Despite being difficult to spot, the *Leuc. plasmidifera* plastid is less reduced than plastids of some other non-photosynthetic taxa characterized by the absence of a genome, exemplified by the chrysophyte genus *Paraphysomonas* (Dorrell et al. 2019). Our investigations indicate that the *raison d’être* for the plastid genome in *Leuc. plasmidifera* (i.e. disregarding its role in the genome maintenance and gene expression itself) is to provide two specific products required for the plastid functions: the (highly simplified) ClpC protein presumably involved in protein turnover in the organelle, and Glu-tRNA serving as an input of a haem biosynthesis pathway housed by the plastid. We identified two additional processes taking place in the organelle and potentially functionally important for the cell as a whole: biosynthesis of a tetrahydrofolate precursor and of tocotrienols. It is possible that the *Leuc. plasmidifera* plastid takes part in additional processes that reach beyond the maintenance of the plastid itself. A prerequisite for a more comprehensive understanding of the physiological roles of the *Leuc. plasmidifera* cryptic plastid is to overcome limitations of the approach taken in this study, i.e. solely bioinformatic reconstructions based on a transcriptome assembly (unlikely to cover all plastid-targeted proteins and frequently not allowing for correct inference of the N-terminal sequences of the proteins, interfering thus with predictions of subcellular targeting). Thus, our future aim is to obtain a high-quality assembled and annotated genome sequence of *Leuc. plasmidifera*, which will provide an excellent resource for addressing diverse aspects of the plastid biology of this organism. Beyond the metabolic pathways themselves, the plastidial features to be investigated should include the repertoire of metabolite transporters mediating the communication of the plastid with the rest of the cell, or the molecular machinery involved in the genome maintenance and gene expression, which seems to exhibit an extreme degree of modification compared to other plastids. Employing biochemical methods may be ultimately needed to experimentally verify at least some of the bioinformatic predictions, such as the production of tocotrienols by the *Leuc. plasmidifera* plastid.

As an unexpected ramification of our work, the plastid is not the only organelle of *Leuc. plasmidifera* that should attract further attention. The features of the long insertion found in one of the mitochondrial *cob* gene introns and of the unusually large circular mitochondrial plasmid related to the insertion seem to point to presently unknown and potentially unique functional aspects of the *Leuc. plasmidifera* mitochondrion. One of the puzzling attributes of both the insertion and the plasmid is the presence of the non-standard tRNA genes: are they used for translation, and if yes, what are the amino acids they are charged with, which codon(s) they decode, and why they are used as all? Similarly enigmatic is the significance of the expanded family of proteins encoded both by the insertion and the plasmid sharing a novel conserved domain lacking discernible homologs in other organisms. The putative occurrence of multiple different members of the family in the *Leuc. plasmidifera* mitochondrion may, speculatively, mean that the proteins form heterooligomers, presumably in the inner mitochondrial membrane (given the fact the conserved N-terminal domain is predicted to include a transmembrane helix). The (nearly) absolute conservation of particular histidine and cysteine residues is notable and suggestive of a specific functionally critical role, such as mediating catalysis of an unknown biochemical reaction, binding of a cofactor or a prosthetic group, or – in the case of the cysteine residue – formation of intermolecular disulfide bonds.

At any rate, the identification of this new protein family indicates that a unique molecular process of unknown physiological significance takes place in the *Leuc. plasmidifera* mitochondrion. Further hints towards possible answers to these questions might be provided by future exploration of metagenomes detected here to include sequences from *Leuc. plasmidifera* relatives. Recovering organellar genomes of these organisms would tell us whether the unusual features of the *Leuc. plasmidifera* plastid and mitochondrial genomes are shared by a broader organismal group, while identification of plasmids related to pLPM would allow us to illuminate the origin of the plasmid and to hopefully also gain some functional insights.

## 4. Taxonomic summary

### Stramenopiles

#### Ochrophyta incertae sedis

##### Leucomyxales Barcytė & M.Eliáš, ord. nov

Description: Heterotrophic unicellular amoeboid eukaryotes with a non-photosynthetic plastid that according to phylogenies inferred from the sequences of the 18S rRNA gene constitute a unique evolutionary lineage typified by *Leucomyxa plasmidifera* Barcytė & M.Eliáš and most closely related to, but separate from, the lineages corresponding to the orders Synchromales S.Horn & Ehlers (including the genus *Synchroma* R.Schnetter) and Chlamydomyxales Archer (including the genus *Chlamydomyxa* W.Archer).

Note: presently with a single family (Leucomyxaceae Barcytė & M.Eliáš), a single genus (*Leucomyxa* Barcytė & M.Eliáš), and a single species (*Leucomyxa plasmidifera* Barcytė & M.Eliáš) described below. The existence of a high diversity of undescribed and uncultured members of the order is indicated by eDNA sequence data.

##### Leucomyxaceae Barcytė & M.Eliáš, fam. nov

Description: Amoeboid heterotrotrophic bacteriovorous protists, with cells containing non- photosynthetic plastids with a genome, thriving primarily in organic-rich terrestrial environments.

Type genus: *Leucomyxa* Barcytė & M.Eliáš

##### *Leucomyxa* Barcytė & M.Eliáš, gen. nov

Description: Amoeboid cells bearing pseudopodia of different types, including anastomosing reticulopodia. Uninucleate cells or multinucleate plasmodia. Highly vacuolated. Flagellate stages present. Cysts formed. Free-living, bacteriovorous, predominantly in soil and other organic-rich terrestrial habitats. Mitochondria with tubular cristae. Reduced non- photosynthetic plastids with a genome retained.

##### Type species (described here): *Leucomyxa plasmidifera*

Etymology: from the Ancient Greek words λευκός (“white”; in reference to the absence of photosynthetic pigments, also points to the presence of a non-photosynthetic plastid, i.e. e leucoplast) and μύξα (“slime”; frequently used as part of generic names of amoeboid protists, such as the related *Chlamydomyxa*).

Note: the genus potentially includes additional species related to the type species, whose existence is presently documented by eDNA sequence data.

##### *Leucomyxa plasmidifera* Barcytė & M.Eliáš sp. nov. (figure 1A–N)

Description: Cells naked, amoeboid with lobopodia and filopodia, or in the form of a multinucleate meroplasmodium (with one to six reticulopodia radiating from a single cell). Cell bodies roundish, elongated or spindle-shaped, stretching 1.5–15 µm in length. Flagellated cells of two types: biflagellate with flagella of unequal length and inserted subapically, cells ovoid to bean-shaped, 3.0 µm long and 2.5 µm wide; and uniflagellate with the single flagellum inserted apically, cells ovoid, 3.0 µm long and 2.0 µm wide. Multiple vacuoles present. Cysts spherical and double-walled, without protrusions, ranging from 2.5–7.0 in diameter.

Holotype designated here: Metabolically inert (cryopreserved) strain ATCC® PRA-24^TM^. Type sequence: rRNA operon sequence deposited at GenBank with the accession number #####.

Type strain: ATCC® PRA-24^TM^ in American Type Culture Collection (www.atcc.org).

Type locality: salt marsh, Virginia, USA.

Etymology: the species epithet refers to the presence of a mitochondrial plasmid in this organism.

## 5. **Material and Methods**

### 5.1. Cultivation and microscopy

*Leukarachnion* sp. PRA-24, further referred to as *Leuc. plasmidifera*, was obtained from The American Type Culture Collection (ATCC®; www.atcc.org). The strain was cultivated in Thermo Scientific^TM^ Biolite 25 cm^2^ cell culture flasks with vented lids (Thermo Fisher Scientific) in seawater cereal grass medium: cereal grass (2,5 g/l), NaCl (24,72 g/l), KCl (0,68 g/l), CaCl2.2H2O (1,36 g/l), MgCl2.6H2O (4,66 g/l), MgSO4.7H2O (6,29 g/l), NaHCO3 (0,18 g/l). The cultures were kept in dark at room temperature and they contained bacteria that served as a food source for *Leuc. plasmidifera* cells. The cultures of *Leuc. plasmidifera* were examined using an Olympus BX53 (Olympus, Tokyo, Japan) fluorescence microscope employing differential interference contrast (DIC). Micrographs were captured using an Olympus DP73 digital camera. Cells were measured using an Olympus micro imagining software cellSens v1.6. For the transmission electron microscopy (TEM) observations, the cells were prepared using high-pressure freezing (HPF). Centrifuged cells were immersed in a cryoprotectant (20% BSA in cell medium) and immediately frozen with a Leica HPM100 high-pressure freezer (Leica Microsystems, Vienna, Austria). The cells were freeze substituted in 2% OsO4 and 100% acetone using an automatic freeze substitution system Leica EM AFS2. The sample was then washed three times in 100% acetone followed by a gradual embedding in EMbed-812 resin at room temperature (resin:acetone – 1:2, 1:1, and 2:1 for one hour per each) with a final overnight embedding in 100% resin. Finally, it was embedded into a fresh degassed resin and polymerized at 60°C for 48 hours. Thin sections were cut on a Reichert-Jung Ultracut E ultramicrotome (Reichert-Jung, Vienna, Austria) and stained using uranyl acetate and lead citrate. Sections were examined and photographed using a JEOL JEM-1011 (JEOL, Tokyo, Japan) electron microscope, equipped with a Veleta camera and the iTEM 5.1 software (Olympus Soft Imaging Solution GmbH, Münster, Germany).

### 5.2. **RNA and DNA extraction**

For RNA extraction, 100 ml of the culture was used, after spinning at 1,500 g for 15 min, the pellet was resuspended in 1ml of TRI Reagent® (Genbiotech), incubated for 5 min, centrifuged at 12,000 g for 15 min on 4 °C and the supernatant was transferred to a fresh tube. After adding of 200 µl of chloroform, shaking, and 15 min of incubation at room temperature, the sample was spun at 12,000 g for 15 min on 4 °C and the aqueous phase was transferred to a fresh tube. It was precipitated with 500 µl of isopropanol, incubated for 30 min on -80 °C, then spun at 12000 g for 8 min on 4 °C, the supernatant was discarded, the pellet washed with 70 % ethanol, dried and dissolved in 50 µl of RNase free water on 55 °C for 15 min. The concentration was checked on NanoDrop Lite Spectrophotometer (Thermo Scientific), after DNase treatment (Thermo Scientific), the concentration was checked on NanoDrop Lite Spectrophotometer again and the quality of RNA was checked by gel electrophoresis (0.8 % (w/v) agarose gel stained with ethidium bromide) using the Gene Ruler 1 kb DNA Ladder (Fermentas) as a standard. For DNA extraction, approximately 200 ml of liquid culture was used. It was centrifuged at 1,500 g for 25 min, the supernatant was discarded and the pellet used for DNA extraction by modified protocol originally used for plants (Dellaporta et al. 1983). Modifications included additional steps of RNAse (RNAse H; the final concentration 0.1 mg/ml) and proteinase treatments (proteinase K; the final concentration 0.2 mg/ml) followed by phenol-chloroform extraction before the final DNA precipitation. The quality and concentration of the isolated DNA quality were checked by gel electrophoresis as for RNA (see above).

### 5.3. **Transcriptome and genome sequencing and assembly**

The transcriptome was sequenced using Illumina HiSeq 2000 2x150bp platform from cDNA libraries prepared using TruSeq RNA sample prep kit v2 (Illumina, San Diego, CA) at Macrogen Inc. (Seoul, South Korea). The raw reads were error-corrected by Rcorrector (Song and Florea 2015), trimmed with Trimmomatic v. 0.32, and assembled into contigs by Trinity assembler v. 2.8.4. (Haas et al. 2013). Protein sequences were then predicted with TransDecoder (https://github.com/TransDecoder/). DNA library was prepared using TruSeq DNA PCR-Free Protocol (Illumina, San Diego, CA) at Macrogen Inc. (Seoul, South Korea), where it was also sequenced on Illumina HiSeq 2000 2x150bp platform. The total amount of 57,000,000 raw reads were trimmed by Trimmomatic v. 0.32. (Bolger et al. 2014). An initial (meta)genome assembly was built with Spades 3.13.0 (Bankevich et al. 2012) and filtered from bacterial contamination as follows: all contigs > 100,000 bp were investigated by BLAST (Altschul et al. 1997) to determine bacterial contamination; using Burrows-Wheeler Aligner (BWA; Li and Durbin 2010) and SAMtools (Li et al. 2009), reads aligned with identified bacterial contigs were removed, surviving reads were reassembled and decontamination procedure was repeated. The final set of surviving reads (7,400,000 reads) was assembled by Spades 3.13.0 using user defined k-mer sizes (-k 21,33,55,77) and “— careful” option to minimize number of mismatches in the final contig. Such a way, we obtained relatively clean genome assembly containing 11,440 contigs > 1,000 bp, three of them identified as fragments of a ptDNA based on TBLASTN searches with common plastid proteins. Subsequent iterative searches of the raw Illumina reads enabled us to unambiguously extend the ptDNA fragments to obtain a full circular-mapping genome sequence including two copies of the inverted repeat typical for plastid genomes. A single gapless scaffold representing the mitochondrial genome was identified with TBLASTN and reference mitochondrial proteins in the initial (meta)genome assembly; the identity of the 3’ region of the scaffold to the 5’ region indicated it corresponds to a complete circular-mapping genome. In the assembly based on filtered reads the mitogenome sequence was incomplete and interrupted in the region corresponding to the 23S rRNA gene, presumably because of the removal of sequencing reads from that region that mapped to the homologous conserved regions in bacterial genomes. The final assemblies of organellar genomes were verified by visual inspection (using Tablet; Milne et al. 2016) of sequencing reads mapped onto the assembled genome sequences with BWA. To investigate the discrepancy between the assembled sequences of the PHD1 transcript and the corresponding genomic region, RNAseq reads were mapped onto the transcriptome assembly by using HISAT2 (Kim et al. 2019) and variability of the read sequences along the PHD1 transcript was inspected in Tablet.

### 5.4. **Annotation of organellar genomes and the mitochondrial plasmid**

The assembled plastid and mitochondrial genomes of *Leuc. plasmidifera* were initially annotated with MFannot (Lang et al. 2023) and the annotation provided by the program was carefully manually checked to ensure that the genes and coding sequences are correctly delimited and that no gene present in the genome is missed. One such extra gene, the short and poorly conserved *tatA*, was identified in the mitogenome and integrated into the annotation. The identity of tRNA genes was additionally checked using tRNAscan-SE v. 2.0 (Chan and Lowe 2019) to correctly annotate initiator and elongator tRNA(cau) species and the lysidinylated Ile-tRNA(cau) cognate to the AUA codon. Varna v3.91 (Darty et al 2009) was used for visualization of predicted secondary structures of tRNAs of specific interest.

Genes for 5S rRNA were searched in both organellar genomes by employing Infernal 1.1.4 (Nawrocki and Eddy 2013) using the covariance models reported by Valach et al. (2014). While no 5S rRNA gene was found in the plastid genome with any of the alternative models employed, the mitochondrial 5S rRNA gene was detected with the model 5S-mito- derived.cm. Further attempts to identify a possible 5S rRNA gene was based on searching for a sequence segment in the unannotated region downstream of the 23S rRNA gene that would fold into an RNA secondary structure characteristic for 5S rRNA. The RNAfold WebServer (http://rna.tbi.univie.ac.at/cgi-bin/RNAWebSuite/RNAfold.cgi) was fed with various subsections of the aforementioned region and the predicted structures were inspected to detect possible structural resemblance to 5S rRNA, including its circularly permuted variants (Valach et al. 2014). Possible homology of the several unidentified ORFs to proteins from other eukaryotes were sought using BLAST, HMMER 3.0 (Eddy 2011), HHpred (Zimmermann et al. 2018), and Phyre2 (Kelley et al. 2015). In addition to the NCBI nr protein sequence database, the EukProt3 database (Richter et al. 2022) containing sequences inferred from genome and transcriptome assemblies of phylogenetically diverse eukaryotes (especially protists) was exploited. For protein products of those ORFs that eluded identification by the aforementioned tools, we built tertiary structure models by using AlphaFold2 (Jumper et al. 021) and compared them with FoldSeek (van Kempen et al. 2023) with default settings against the vast collection of experimentally determined and AlphaFold-predicted protein structural models offered for searching by the FoldSeek server (https://search.foldseek.com/search). The presence of transmembrane domains in plastid-encoded proteins was analysed using TMHMM server v. 2.0 (http://www.cbs.dtu.dk/services/TMHMM/). The plastid genome map was obtained using OGDRAW v.1.3.1 (https://chlorobox.mpimp-golm.mpg.de/OGDraw.html; Greiner et al. 2019). Data on the gene presence/absence in plastomes of various ochrophytes were adopted from Barcytė et al. (2021) and updated by adding details from our *Leuc. plasmidifera* plastome annotation.

### 5.5. **Analysis of nucleus-encoded plastid-targeted proteins**

Homologs of proteins of specific interest encoded by the *Leuc. plasmidifera* nuclear genome were searched by different complementary approaches. TBLASTN was employed to search the transcriptome assembly for homologs of well-characterised reference plastid proteins gathered from databases (especially KEGG; https://www.genome.jp/kegg/pathway.html) or the literature. HMMER was employed as a more sensitive tool to search against the database of *Leuc. plasmidifera* protein sequences inferred from the transcriptome assembly. Hidden Markov models (HMMs) were built from seed alignments of appropriate protein families or domains downloaded from the database Pfam (http://pfam.xfam.org/; Mistry et al. 2021) or from custom protein sequence alignments (obtained by using MAFFT version 7; https://mafft.cbrc.jp/alignment/server/; Katoh and Standley 2013) and used as queries for the HMMER searches. Hits with significantly low e-values (typically <0.001) were evaluated by BLASTX or BLASTP searches against the NCBI nr protein sequence database to remove false hits, bacterial contaminants (most frequently encountered were contaminants from a *Vibrio* sp.), or sequences representing a different member (paralog or pseudoparalog) of the respective protein family than the protein searched for. The presence of an N-terminal signal peptide, a prerequisite for the protein to be potentially plastid targeted (see section 3.4.) in the candidates was evaluated by using TargetP 2.0 (“non-plant” setting; https://services.healthtech.dtu.dk/service.php?TargetP-2.0; Almagro Armenteros et al. 2019), Predotar (https://urgi.versailles.inra.fr/predotar/; Small et al. 2004), PredSL (set to “non-plant sequences”; http://aias.biol.uoa.gr/PredSL/input.html; Petsalaki et al. 2006), PrediSi (http://www.predisi.de/; Hiller et al. 2004), and DeepLoc 2.0 (Thumuluri et al. 2022). The former three tools provided also prediction of possible mitochondrial targeting of the proteins, which was additionally tested using MitoFates (arbitrarily with the default setting to “fungi”; http://mitf.cbrc.jp/MitoFates/cgi-bin/top.cgi; Fukasawa et al. 2015). The presence of Fe-S clusters in proteins was predicted by using MetalPredator Valasatava et al. 2016).

### 5.6. **Phylogenetic analyses**

The sequence of the 18S rRNA gene of *Leuc. plasmidifera* PRA-24 was extracted from a scaffold in the genome assembly and its confirmed by comparison to the corresponding sequence in the transcriptome assembly. Environmental 18S rRNA gene sequences specifically related to that of *Leuc. plasmidifera* PRA-24 were identified by BLASTN searches against the NCBI nr nucleotide sequence database and against metagenome assemblies in the NCBI WGS database. Sequences with a putative specific relationship to that from *Leuc. plasmidifera* PRA-24 indicated by a preliminary FastTree analysis at the ETE3 server (https://www.genome.jp/tools-bin/ete) were further checked for chimeric assembly by comparing their different segments to the NCBI nr nucleotide sequence database with BLASTN and considering the identity of the best hits. 49 sequences with no evidence for being chimeric were retained and combined with the 18S rRNA gene sequence from *Leuc. plasmidifera* PRA-24 and 53 additional organisms representing all the known major ochrophyte lineages. A multiple sequence alignment was constructed with MAFFT using the L-INS-i method. The alignment was inspected by eye and poorly conserved positions were removed manually. A ML tree was computed with IQ-TREE multicore version 2.2.5 (Minh et al. 2020) using a substitution model chosen by the program as best fitting the data (TIM2+F+I+R5). Branch support was evaluated by non-parametric bootstrapping with 100 replicates. Thirty plastid-genome encoded proteins present in at least one species of the pair of non-photosynthetic ochrophytes “*Spumella*” sp. NIES-1846 (Dorrell et al. 2019) and *Leuc. plasmidifera* and suitable for phylogenetic analysis (marked in supplementary electronic material, table S3) were selected to build a supermatrix for phylogenomic inference. For each protein, orthologs were collected from a set of reference ochrophyte plastid genomes (including representatives of all classes for which plastid genome sequence is available, and including also various non-photosynthetic ochrophytes for comparison) as well as plastid genomes of selected non-ochrophytes (to provide an outgroup); the list of the genomes analysed and the sequence identifiers are provided in supplementary electronic material, table S4. The orthologous protein sequences (in some cases modified compared to the available annotation to fix obvious mistakes in the definition of the coding sequence start or to account for apparent plastid genome sequence assembly errors) were aligned using MAFFT with the E-INS-i option. The alignments were checked visually, trimmed with trimAl (Capella-Gutierrez et al. 2009), and concatenated with FASconCAT (Kück and Meusemann 2010) to yield a supermatrix of 6,627 aligned amino acid positions. The ML tree was inferred using IQ-TREE multicore version 2.0.3 with the substitution model LG+C60+F+G and 100 nonparametric bootstrap replications. A sequence set for the plastid 16S rRNA gene, collected by extracting the respective regions from plastid genome sequences or identified by BLASTN in the NCBI nr database, was aligned and trimmed in the same manner as described for the plastid protein sequence analyses, followed by a separate ML analysis in IQ-TREE employing the GTR+F+R4 substitution model, respectively, and 100 nonparametric bootstrap replications. All phylogenetic trees were visualised using FigTree 1.4.4 (Rambaut 2009) or iTOL (Letunic and Bork 2021) and post-processed in a graphical editor.

## Data Accessibility

The sequences of the 18S rRNA gene, plastid and mitochondrial genomes, and the putative mitochondrial plasmid from *Leucomyxa plasmidifera* PRA-24 were deposited at GenBank with the accession numbers ######-######. Raw RNAseq reads and transcriptome assembly from the same species are available under the NCBI BioProject ########. The (low-coverage and fragmented) draft genome assembly and sequence alignments analysed in the study are provided at Figshare#####.

## Author Contributions

D.B.: conceptualization, formal analysis, investigation, validation, visualization, writing – original draft, writing – review & editing; K.J.: data curation, formal analysis, investigation, methodology, validation, visualization, writing – review & editing; T.P: formal analysis, investigation, resources, writing – review & editing; T.Y.: data curation, investigation, visualization, writing – review & editing; T.Š.: data curation, formal analysis, supervision, validation, writing – review & editing; A.E.: project administration, resources, writing – review & editing; M.E.: conceptualization, data curation, formal analysis, funding acquisition, investigation, methodology, supervision, validation, writing – original draft, writing – review & editing.

## Conflict of interest declaration

We declare we have no competing interests.

## Funding

This work was supported by the Czech Science Foundation (project 17-21409S and 21-19664S) and the project “CePaViP”, supported by the European Regional Development Fund, within the Operational programme for Research, Development and Education (CZ.02.1.01/0.0/0.0/16_019/0000759). TP was supported by the Charles University (project UNCE 204069).

## Supplementary Material

The Supplementary Material for this article can be found online at: xxxxxxxxx.

## Notes

### Competing Interest Statement

The authors have declared no competing interest.

### Summary of Updates

The scope of the manuscript has been been expanded to include an analysis of environmental DNA data, which together with a more thorough taxonomic work on the organism studied resulted in our decision to consider "Leukarachnion sp. PRA-24" a representative of a new species in a new genus, proposed to be named Leucomyxa plasmidifera. We have also added completely new sections reporting on the mitochondrial genome of the organism and a unique mitochondrial plasmid discovered in it. Finally, we have expanded and refined the reconstruction of the metabolic functions of the Leuc. plasmidifera plastid. The new version is supposed to be much more accurate and scientifically robust compared to the originally submitted version.

