## Supplementary notes & figures for "A cryptic plastid and a novel mitochondrial plasmid in *Leucomyxa plasmidifera* gen. and sp. nov. (Ochrophyta) push the frontiers of organellar biology"

**This file contains supplementary material to the paper (except for supplementary tables included in a separate xlsx file):**

**Supplementary note S1.** Additional details on the ultrastructure of *Leucomyxa plasmidifera* PRA-24

**Supplementary note S2.** Additional discussion on the relationship of *Leucomnyxa plasmidifera* PRA-24 to previously described taxa

**References to supplementary notes**

**Supplementary tree data:** A full version (in the Newick format) of the 18S rRNA gene tree presented in the main text as figure 2.

**Figure S1.** Ultrastructure of *Leucomyxa plasmidifera* PRA-24.

**Figure S2.** Assessment of the representativeness of the *Leucomyxa plasmidifera* PRA-24 transcriptome assembly using BUSCO.

**Figure S3.** The meaning of codons in protein-coding genes in the organellar genomes of *Leucomyxa plasmidifera* PRA-24.

**Figure S4.** Maximum likelihood tree inferred from plastidial 16S rRNA sequences.

**Figure S5.** Two versions of the PHD1 gene in *Leucomyxa plasmidifera* PRA-24.

**Figure S6.** Mutations of catalytically critical residues in the TrpF-TrpC fusion protein from *Leucomyxa plasmidifera* PRA-24.

**Figure S7.** Unusual tRNAs specified by the *Leucomyxa plasmidifera* PRA-24 mitochondrial genome and its novel mitochondrial plasmid.

**Figure S8.** Novel protein family encoded by the *Leucomyxa plasmidifera* PRA-24 mitochondrial genome and its novel mitochondrial plasmid.

**Supplementary note S1.** Additional details on the ultrastructure of *Leucomyxa plasmidifera* PRA-24

We used a protocol employing high-pressure freezing (HPF), which is supposed to secure a better preservation of membranes compared to the classical chemical fixation (Studer et al. 1989). We observed three life forms of *Leuc. plasmidifera* in our TEM specimens: the vegetative cells (figure S1A), cysts (figure S1B), and flagellates (figure S1G). While vegetative cells did not possess any prominent cell wall above the plasma membrane, cysts were covered by a thick double-layered wall (figure S1B) which thickness varied among cells, probably depending on the cyst age. The outer layer of the cyst wall was fibrous, with fibres occasionally projecting outward and forming a fuzzy surface (figure S1C). The inner layer was more electron dense and of uneven thickness depending on cell membrane invaginations (figure S1B). The vegetative cells typically possessed one to three nuclei of variable position. Mitochondria were highly abundant and contained tubular cristae (figure S1D). The cytoplasm was dominated by ribosomes and vacuoles bounded by a single membrane. No classical stacked Golgi body (dictyosome) was observed, which is in agreement with the previous investigation (Grant et al. 2009). Multivesicular bodies were detected (figure S1A), as well as different electron-dense pleomorphic vesicles or bodies of an unclear nature (figure S1E,F), and cytoplasmic microtubules (figure S1A,H).

Of the highest interest was to identify putative plastids in *Leuc. plasmidifera*, which, in analogy with other ochrophytes, we expected to be located close to the nucleus and having the outermost membrane continuous with the nuclear envelope. Such structures, interpreted as leucoplasts, were detected in some other non-photosynthetic ochrophytes, including the dictyochophyte *Pteridomonas danica* (Sekiguchi et al. 2002) or species of the chrysophyte genus *Spumella* (Jeong et al. 2021). A notable example is also the four membrane-containing apicoplast of the apicomplexan parasite genus *Plasmodium* detected in the close proximity to the nucleus despite the lack of continuity between the outermost membranes of the two organelles (Lemgruber et al. 2013). A membranous structure bounded by potentially more than two membranes and position close to the nucleus was also observed once in our TEM preparations of *Leuc. plasmidifera* cells (figure S1F), but we are reluctant to conclude this is the sought-after plastid. Noteworthy, in a *Leuc. plasmidifera* relative, the amoeboid photosynthetic alga *Chlamydomyxa montana*, no ER membranes were evident under the TEM investigation (Pearlmutter and Timpano 1984), though the presence of the two membranes of the periplastidial endoplasmic reticulum was confirmed in another *Chlamydomyxa* species, *C. labyrinthuloides* (Wenderoth et al. 1999). Another *Leuc. plasmidifera* relative, namely the genus *Synchroma*, displays peculiar plastid arrangements termed "plastid complexes" (Horn et al. 2007), although these are not found in other organisms in the same clade (*Guanchochroma* and *Chrysopodocystis*; Schmidt et al. 2015). The specific "search image" of the elusive *Leuc. plasmidifera* plastid is thus not obvious *a priori*, and as the cryptic plastid most likely lacks thylakoids, it may be very difficult to tell it apart from other membranous compartments in the *Leuc. plasmidifera* cells without employing a target-specific approach, such as immunogold labelling.

**Supplementary note S2.** Additional discussion on the relationship of *Leucomnyxa plasmidifera* PRA-24 to previously described taxa

Grant et al. (2009) speculated about a close relationship of the strain PRA-24 to *Leukarachnion batrachospermi*, but Geitler's account on that species, very detailed and meticulous, does not provide a strong case for such a conjecture (Geitler 1942). He found *Leuk. batrachospermi* once in a freshwater stream (Lunzer Sebach, Austria) as a parasite of multicellular red algae then assigned to the genus *Batrachospermum*, thriving in the mucilage produced by these algae. This amoeboid organism was capable of forming large (up to 2 mm broad) highly reticulated plasmodia (meroplasmodia) and made living by seeking out spermatangia and carposporangia of the host red algae, embracing them, and removing reproductive cells to digest them in large digestive vacuoles. Geitler noted that apart from the reproductive *Batrachospermum* cells, the plasmodium generally did not contain other food bodies, with very few cases of putative bacteria in digestive vacuoles observed. Geitler mentioned two *Batrachospermum* species as the *Leuk. batrachospermi* hosts, one unidentified and the other considered to be *Batrachospermum boryanum* (now known as *Sheathia boryana*; Salomaki et al. 2014), and he indicated a high incidence of infection by *Leuk. batrachospermi*, with not a single uninfected thallus found among hundreds examined. In contrast, he reported that no *Leuk. batrachospermi* was found in association with other algae and cyanobacteria in the same locality, further attesting to this organism being a highly specialised in terms of its occurrence, behaviour and diet. Geitler later documented a similar, "*Leukarachnion*-like" organism, found as a fixed specimen on microscopic slides exposed for several weeks to running water in another Austrian stream, but its behaviour and nutrition mode could not be established for a lack of living material, leaving its relationship to *Leuk. batrachospermi* uncertain (Geitler 1958/1959).

The hypothesis by Grant et al. (2009) about a specific relationship of the PRA-24 strain to *Leuk. batrachospermi* implies the latter organism is a stramenopile. However, Geitler (1942) himself noted that the feeding specialization of *Leuk. batrachospermi* indicates relationship to Vampyrellida, a generally eukaryovorous and predominantly algivorous group of amoeboid protists, now known to belong to Rhizaria, which employ different feeding strategies including not only the spectacular eponymous "sucking" of the cell content but also the more conventional phagocytic engulfment of whole cell (Hess and Suthaus 2022), like seen in *Leuk. batrachospermi*. Furthermore, some vampyrellids form large reticulated plasmodia not unlike *Leuk. batrachospermi*, and the vampyrellid life cycle includes thick-walled resting stages (spores) comparable to the cysts described for *Leuk. batrachospermi* yet lack flagellated stages, again in accord with Geitler's report. Notably, eDNA data revealed the existence of a number of lineages up to the family level coming from freshwater habitats and representing vampyrellid that are yet to be described or assigned to previously described taxa (Hess and Suthaus 2022); it is possible that *Leuk. batrachospermi* matches one of these lineages.

Having discounted the relationship of the PRA-24 strain to *Leuk. batrachospermi*, we considered additional previously described meroplasmodial heterotrophic protists as possible PRA-24 relatives. An extensive list of such protists was assembled by Berney et al. (2015), but of those found in terrestrial and freshwater environments all are known or suspected to belong among vamyrellids or leptomyxid amoebozoans. Our literature survey revealed one other previously described heterotrophic plasmodial reticulopodial protists, *Leukapsis vorax* described by Pascher (1940) and not studied since then. However, as concluded by Pascher himself, this organism is most likely a non-photosynthetic chrysophyte, considering the characteristic details of the formation and morphology of the cysts. Thus, it seems that the PRA-24 strain does not match any of the previously described genus (let alone species), justifying our decision to formally describe it as a new species in a new genus, *Leucomyxa plasmidifera* gen. et sp. nov.

### References to supplementary notes

Berney C, Geisen S, Van Wichelen J, Nitsche F, Vanormelingen P, Bonkowski M, Bass D. 2015 Expansion of the 'reticulosphere': diversity of novel branching and network-forming amoebae helps to define Variosea (Amoebozoa). *Protist* **166**, 271–295. (doi:10.1016/j.protis.2015.04.001)

Geitler L.1942 Ein neue filarplasmodialer Organismus, *Leukarachnion batrachospermi*, und seine Lebensweise. *Biol. Zentralbl.* **62**, 541–549.

Geitler L. 1958/1959 Über einen leukarachnion-ähnlichen filarplasmodialen rhizopoden. *Arch. Protistenk.* **103**, 573–580.

Grant J, Tekle YI, Anderson OR, Patterson DJ, Katz LA. 2009 Multigene evidence for the placement of a heterotrophic amoeboid lineage *Leukarachnion* sp. among photosynthetic stramenopiles. *Protist* **160**, 376–385. (doi:10.1016/j.protis.2009.01.001)

Hess S, Suthaus A. 2022 The vampyrellid amoebae (Vampyrellida, Rhizaria). *Protist* **173**, 125854. (doi:10.1016/j.protis.2021.125854)

Horn S, Ehlers K, Fritzsch G, Gil-Rodríguez MC, Wilhelm C, Schnetter R. 2007 *Synchroma grande* spec. nov. (Synchromophyceae class. nov., Heterokontophyta): An amoeboid marine alga with unique plastid complexes. *Protist* **158**, 277–293. (doi:10.1016/j.protis.2007.02.004)

Jeong M, Kim JI, Nam SW, Shin W. 2021 Molecular phylogeny and taxonomy of the genus *Spumella* (Chrysophyceae) based on morphological and molecular evidence. *Front. Plant Sci.* **12**, 758067. (doi:10.3389/fpls.2021.758067)

Lemgruber L, Kudryashev M, Dekiwadia C, Riglar DT, Baum J, Stahlberg H, Ralph SA, Frischknecht F. 2013 Cryo-electron tomography reveals four-membrane architecture of the *Plasmodium* apicoplast. *Malar. J.* **12**, 25. (doi:10.1186/1475-2875-12-25)

Pascher A. 1940 Filarplasmodiale Ausbildungen bei Algen. *Arch. Protistenkn.* **94**, 295–309.

Pearlmutter NL, Timpano P. 1984 The biology of *Chlamydomyxa montana*: Ultrastructure of the cyst. *Protoplasma* **122**, 68–74. (doi:10.1007/BF01279438)

Schmidt M, Horn S, Ehlers K, Wilhelm C, Schnetter R. 2015 *Guanchochroma wildpretii* gen. et spec. nov. (Ochrophyta) provides new insights into the diversification and evolution of the algal class synchromophyceae. *PLoS One* **10**, e0131821. (doi:10.1371/journal.pone.0131821)

Sekiguchi H, Moriya M, Nakayama T, Inouye I. 2002 Vestigial chloroplasts in heterotrophic stramenopiles *Pteridomonas danica* and *Ciliophrys infusionum* (Dictyochophyceae). *Protist* **153,** 157–167. (doi:10.1078/1434-4610-00094)

Studer D, Michel M, Müller M. 1989. High pressure freezing comes of age. *Scanning Microsc. Suppl.* **3**, 253–268.

Wenderoth K, Marquardt J, Fraunholz M, Van de Peer Y, Wastl J, Maier UG. 1999 The taxonomic position of *Chlamydomyxa labyrinthuloides*. *Eur. J. Phycol.* **34**, 97–108. (doi:10.1080/09670269910001736152)

**Supplementary tree data:** A full version (in the Newick format) of the 18S rRNA gene tree presented in the main text as figure 2.

(Leucomyxa_plasmidifera_gen._et_sp._nov.:0.0000011419,((CAAACL010049881.1_1251-2364_bioreactor_metagenome_genome_assembly_NODE_49881_length_2364_cov_43.645734:0.0000011419,(CAAABU010155282.1_c562-1_bioreactor_metagenome_genome_assembly_NODE_155282_length_562_cov_14.065089:0.0000011419,((((((((((PJTQ01003331.1_898-2626_Soil_metagenome_Ga0160482_103331:0.0000028613,MW433495.1_Uncultured_eukaryote_clone_ASV221:0.0000011419)77:0.0017687305,MW433508.1_Uncultured_eukaryote_clone_ASV234_small_subunit_ribosomal_RNA_gene__partial_sequence:0.0016648077)56:0.0035632780,(((((((((OBEP011496425.1_metagenome_genome_assembly_contig__so01A-GAAACAATAATCAGCGCGGA-contig-1-11697:0.0000011419,FPLL01004553.1_metagenome_genome_assembly_contig__4553:0.0000028448)100:0.0016458901,(OBEP011514510.1_metagenome_genome_assembly_contig_so01B-ATTATTCTATTTGTACCCGA-contig-1-11308:0.0000011419,FPLS01014300.1_metagenome_genome_assembly_contig__14300:0.0007723203)98:0.0014805785)76:0.0011553235,(JAANMX010006862.1_c5963-4841_Bioreactor_metagenome_contig-100_6862:0.0000024879,(((UWTO01006122.1_2981-4705_bioreactor_metagenome_genome_assembly_NODE_6122_length_7645_cov_39.288406:0.0000032643,UWTF01162373.1_bioreactor_metagenome_genome_assembly_NODE_162373_length_563_cov_20.761811:0.0000028639)85:0.0021908805,KU658997.1_Uncultured_eukaryote:0.0183418488)17:0.0025984041,KU820728.1_Uncultured_eukaryote_clone_OTU_98:0.0017840368)17:0.0000025353)21:0.0046323000)29:0.0036427412,KF357520.1_Uncultured_chrysophyte_clone_G8:0.0059054702)47:0.0025578832,((JAITZQ014503998.1_c1196-1_Soil_metagenome_SD2897-2912_k127_8301772:0.0062168923,JAITZO010010874.1_c3380-1655_Soil_metagenome_SD2911_NODE_10874_length_3635_cov_0.921608:0.0000011419)100:0.0093850206,(((OQ786765.1_Chlamydomyxa_labyrinthuloides_isolate_FFRP_1__Free-form_:0.0353186340,(KF443035.1_Chlamydomyxa_labyrinthuloides_strain_P42150__CCAM_:0.0006582810,AJ130893.1_Chlamydomyxa_labyrinthuloides:0.0094206475)100:0.0505972498)100:0.0295083643,((KF925343.1_Synchroma_pusillum_CCMP3072:0.0216346741,DQ788730.1_Synchroma_grande_CCMP2876:0.0244766502)39:0.0013215459,(KF443034.1_Guanchochroma_wildpretii:0.1268326472,KF443036.1_Chrysopodocystis_socialis_AC38:0.1508610369)100:0.1992082674)100:0.0571985508)97:0.0194084988,((((U73221.1_Synura_sphagnicola:0.0573839754,JX946337.1_Mallomonas_papillosa_DMJMpa2:0.0197999777)75:0.0095478291,((FM955256.1_Hydrurus_foetidus:0.0196185843,KY575274.1_Ochromonas_triangulata_strain_A14_651:0.0344300892)29:0.0061261714,(M87332.1_Chromulina_chionophila_CCMP261:0.0481984200,HQ710558.1_Hibberdia_magna_culture_CCMP_453:0.0360264907)46:0.0076514326)38:0.0042945851)30:0.0072897836,AF174376.1_Paraphysomonas_foraminifera:0.0290766802)100:0.0306889997,((((((((MW750343.1_Paralia_sulcata_strain_CNS00428:0.0464049864,HQ912614.1_Stellarima_microtrias_strain_CCMP806:0.1169880953)25:0.0057654482,(((AJ535191.1_Guinardia_flaccida_clone_p788:0.0823794146,AJ535180.1_Corethron_inerme_clone_p534:0.1204093988)83:0.0094505257,OL851898.1_Tenuicylindrus_belgicus_strain_CNS000429:0.1638419986)38:0.0030085191,AY485452.1_Thalassiosira_pseudonana_CCMP1335:0.0958328444)39:0.0040887352)95:0.0215355182,AF167154.1_Triparma_eleuthera_RCC208:0.0640975414)100:0.0354931160,MW045617.1_Olisthodiscus_luteus_K0444:0.1174600375)67:0.0163887268,((HQ007250.1_Trachydiscus_minutus_CCALA_838:0.0745660421,FJ858972.1_Vischeria_punctata_UTEX_153:0.0628738406)95:0.0239996678,(AB183614.1_Polypodochrysis_teissieri_MBIC10541:0.0444750014,AB042204.1_Phaeomonas_parva:0.0531799500)100:0.0542527486)29:0.0048764927)17:0.0044207840,((AY788931.1_Fibrocapsa_japonica_CCMP1661:0.0331993092,(JX026932.1_Haramonas_dimorpha_CCMP2053:0.0273645971,DQ191680.1_Chattonella_subsalsa_CCMP2191:0.0417285729)99:0.0179379180)98:0.0148714658,(AJ579337.1_Botrydiopsis_pyrenoidosa_SAG_3183:0.0325033031,(((MK189081.1_Chrysoparadoxa_australica_EC13:0.0484659629,(((AF083399.1_Heterococcus_caespitosus:0.0354603805,AM490824.1_Tribonema_minus_SAG_880.3:0.0451097778)66:0.0107226541,AM490831.1_Pleurochloridella_botrydiopsis_strain_CCMP1665:0.0611629869)92:0.0179951117,MT582117.1_Nematochrysis_sessilis_var._vectensis_strain_A14_626:0.0928075571)78:0.0171935559)73:0.0087942281,(AJ295822.1_Antarctosaccion_applanatum:0.0697125006,AB365206.1_Tetrasporopsis_fuscescens_SAG20.88:0.0611782516)100:0.0549306153)50:0.0123632497,(((AB365203.1_Phaeothamnion_confervicola_CCMP637:0.0177543639,MK775674.1_Stichogloea_fawleyi_CCMP_2276:0.0389489530)100:0.0282743593,AB365192.1_Aurearena_cruciata:0.1256675321)68:0.0190888371,(MN994274.1_Schizocladia_ischiensis_culture_KU-MACC_KU-333:0.0594285917,((AB087108.1_Dictyota_linearis:0.0368099878,MT582127.1_Ishige_sinicola:0.0290568153)100:0.0342435149,AB252657.1_Discosporangium_mesarthrocarpum:0.0463680298)99:0.0142717115)99:0.0191717322)29:0.0089992311)44:0.0108239027)97:0.0258121334)76:0.0083574804)22:0.0060100448,((U14386.1_Pelagococcus_subviridis_CCMP1429:0.0161071618,HQ710574.1_Aureoumbra_lagunensis_CCMP_1510:0.0443804729)100:0.0674281772,((U14385.1_Octactis_speculum_CCMP1381:0.0657369626,KF422624.1_Florenciella_sp._RCC1587:0.0287327318)95:0.0145700982,((HQ710560.1_Pseudopedinella_elastica_CCMP_716:0.0216563807,AB081640.1_Pteridomonas_danica:0.0493753970)100:0.0261280930,(KF422605.1_Rhizochromulina_cf._marina_CCMP_1243:0.0386499027,L37205.1_Ciliophrys_infusionum:0.0794909824)100:0.0333992643)100:0.0579252640)100:0.0395825007)25:0.0115074424)91:0.0188345455,AF185051.1_Picophagus_flagellatus:0.1007985046)97:0.0190113324)100:0.0246912723)94:0.0169313525)74:0.0092649903)28:0.0045977519,(((LN580544.1_Uncultured_eukaryote_clone_SIBT770_N12D0_18S_E:0.0013997231,(LN575944.1_Uncultured_eukaryote_clone_SIGF1981_N11D1_18S_E:0.0071118959,LN581988.1_Uncultured_eukaryote_clone_SIBH1184_N12D2_18S_E:0.0115583259)61:0.0000032235)18:0.0000024362,LN580474.1_Uncultured_eukaryote_clone_SIBT1361_N12D0_18S_E:0.0000011419)45:0.0000011419,LN581250.1_Uncultured_eukaryote_clone_SIBO712_N12D1_18S_E:0.0028112958)98:0.0355347469)41:0.0027652032,KF188077.1_Uncultured_stramenopile_clone_S5_54:0.0013666876)16:0.0000024612,(JQ480041.1_Uncultured_eukaryote_clone_SIP_OH_B9:0.0000023005,JQ480039.1_Uncultured_eukaryote_clone_SIP_CH_A3:0.0022084096)60:0.0022187247)18:0.0028579326,(KF188080.1_Uncultured_stramenopile_clone_S5_11:0.0068834982,KF188078.1_Uncultured_stramenopile_clone_S5_51:0.0000011419)62:0.0000031434)61:0.0088701444)19:0.0022317090,(JACZCT010346666.1_Soil_metagenome_isolate_MS_1_k53_1501083_1:0.0107891160,LN582247.1_Uncultured_eukaryote_clone_SIBH717_N12D2_18S_E:0.0013791393)11:0.0000013211)0:0.0000016738,LN582141.1_Uncultured_eukaryote_clone_SIBH386_N12D2_18S_E:0.0013805325)0:0.0000028039,LN582102.1_Uncultured_eukaryote_clone_SIBH1443_N12D2_18S_E:0.0013806182)3:0.0000032367,LN581937.1_Uncultured_eukaryote_clone_SIBH1035_N12D2_18S_E:0.0000011419)2:0.0000022753,LN581959.1_Uncultured_eukaryote_clone_SIBH1099_N12D2_18S_E:0.0000011419)31:0.0022566875,((ADGO01002129.1_c380-1_Compost_metagenome_contig02143:0.0000025238,LN585919.1_Uncultured_eukaryote_clone_SIHB.1.722_N9D0_16S_B:0.0000023540)100:0.0096502503,(((LN587345.1_Uncultured_eukaryote_clone_SICY1132_N9D3_18S_E:0.0013705918,((LN585912.1_Uncultured_eukaryote_clone_SIHB.1.691_N9D0_16S_B:0.0000011419,(((LN580971.1_Uncultured_eukaryote_clone_SIBO1040_N12D1_18S_E:0.0013705903,LN575438.1_Uncultured_eukaryote_partial_clone_SIGB572_N11D1_16S_A:0.0018001308)7:0.0000011419,LN576801.1_Uncultured_eukaryote_clone_SICF1146_N11D2_18S_E:0.0000011419)2:0.0000011419,LN587530.1_Uncultured_eukaryote_clone_SICY699_N9D3_18S_E:0.0000011419)2:0.0000011419)3:0.0000011419,LN582942.1_Uncultured_eukaryote_clone_SICZ802_N12D4_16S_A:0.0000011419)5:0.0000011419)13:0.0000032686,((LN583190.1_Uncultured_eukaryote_clone_SIFB530_N12D4_18S_E:0.0155818114,LN583280.1_Uncultured_eukaryote_clone_SIFB844_N12D4_18S_E:0.0301390034)36:0.0013773258,LN583083.1_Uncultured_eukaryote_clone_SIFB1365_N12D4_18S_E:0.0093397565)69:0.0018307370)29:0.0000027015,LN577349.1_Uncultured_eukaryote_clone_SICA1260_N11D3_18S_E:0.0013886042)96:0.0038789871)61:0.0007554843)10:0.0002999462,KU658388.1_Uncultured_eukaryote:0.0000011419)45:0.0000023060)28:0.0000011419)19:0.0000024352,CAAABY010020197.1_c1114-1_bioreactor_metagenome_genome_assembly_NODE_20197_length_2421_cov_14.775993:0.0000011419)43:0.0000011419,EU528037.1_Uncultured_stramenopile_clone_S2_clone_1:0.0000011419);

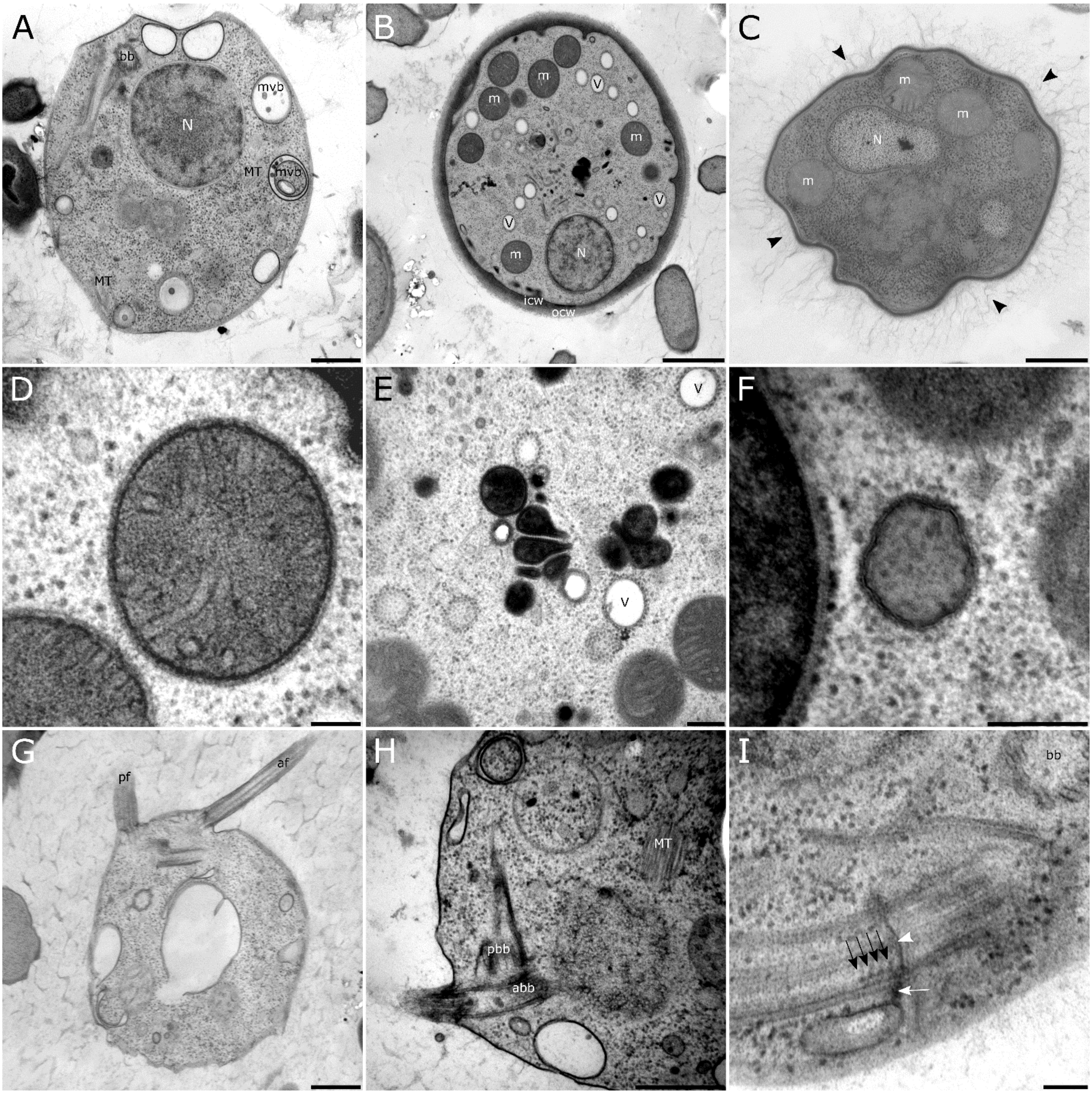


**Figure S1.** Ultrastructure of *Leucomyxa plasmidifera* PRA-24. (A) General organization of the cell (B) Cyst with a thick double-layered cell wall. (C) Fuzzy surface (black arrowheads) of the outer cell wall layer. (D) Mitochondrion with tubular cristae. (E) Aggregation of pleomorphic vesicles with an electron-dense content. (F) Vesicle of an unknown identity. (G) Biflagellate cell. (H) Basal bodies of anterior and posterior flagella. (I) Transition plate (white arrowhead) with four gyres (black arrows) of a transition helix of the anterior flagellum. White arrow points to a dense band connecting axonemal doublets to an unidentified structure. Abbreviations: abb – anterior basal body; af – anterior flagellum; bb – basal body; icw – inner cell wall; ly – lysosome; m – mitochondrion; MT – microtubules; mvb – multivesicular body; N – nucleus; ocw – outer cell wall; pbb – posterior basal body; pf – posterior flagellum; v – vacuole. Scale bars: 0.1 μm (D,H), 0.2 μm (E,I), 0.5 μm (A,C,F,G), 1 μm (B).

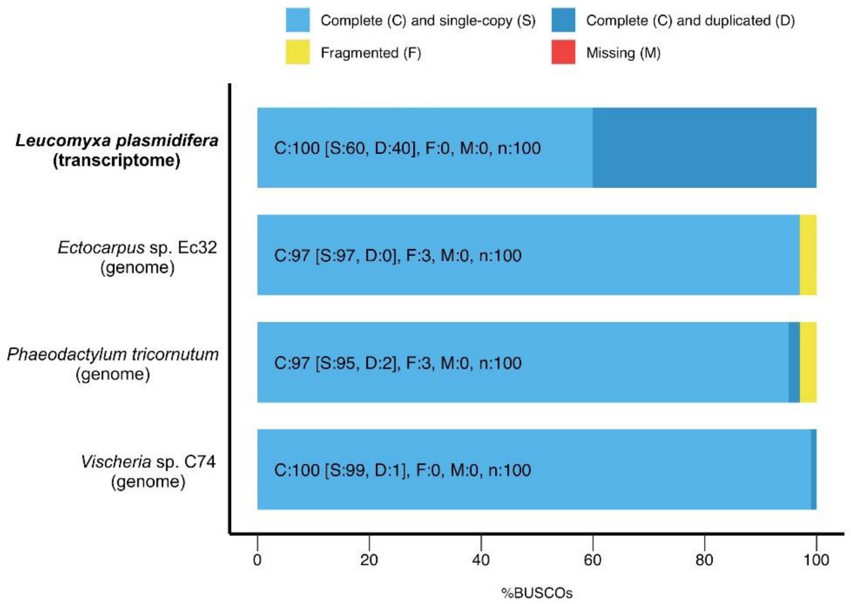


**Figure S2.** Assessment of the representativeness of the *Leucomyxa plasmidifera* PRA-24 transcriptome assembly using BUSCO. The stramenopiles_odb10 reference dataset was used for the analysis. Three different ochrophytes with highly complete genome sequences were included for comparison.

**A**

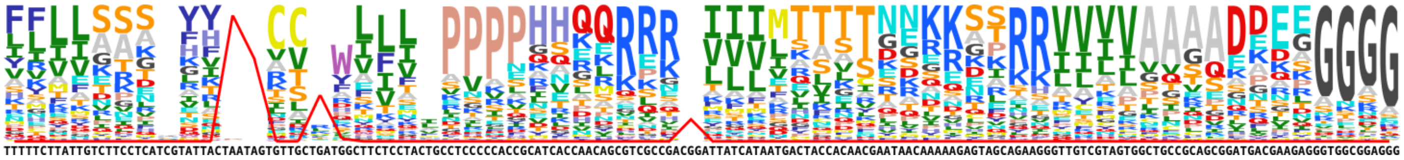

**B**

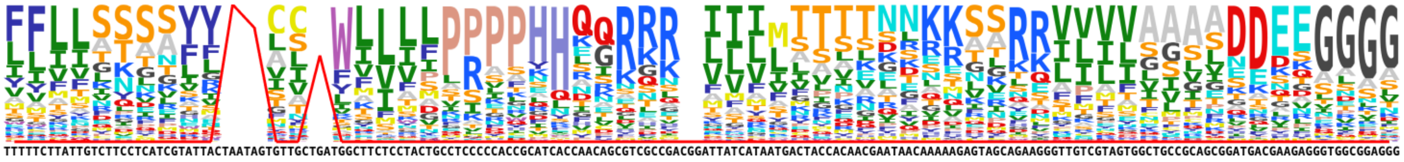

**Figure S3.** The meaning of codons in protein-coding genes in the organellar genomes of *Leucomyxa plasmidifera* PRA-24. The codon meaning in plastid (A) and mitochondrial (B) genes was estimatged with FACIL. The amino acid logo for each codon is inferred from the distribution of the codons at conserved amino acid positions in homologous sequences, with the most likely codon meaning corresponding to the top-most amino acid in the column. The red line indicates the strength of the signal for the codon serving as a termination codon. The plot does not show any signal for a codon reassignment in the *Leuc. plasmidifera* plastid or mitochondrion.

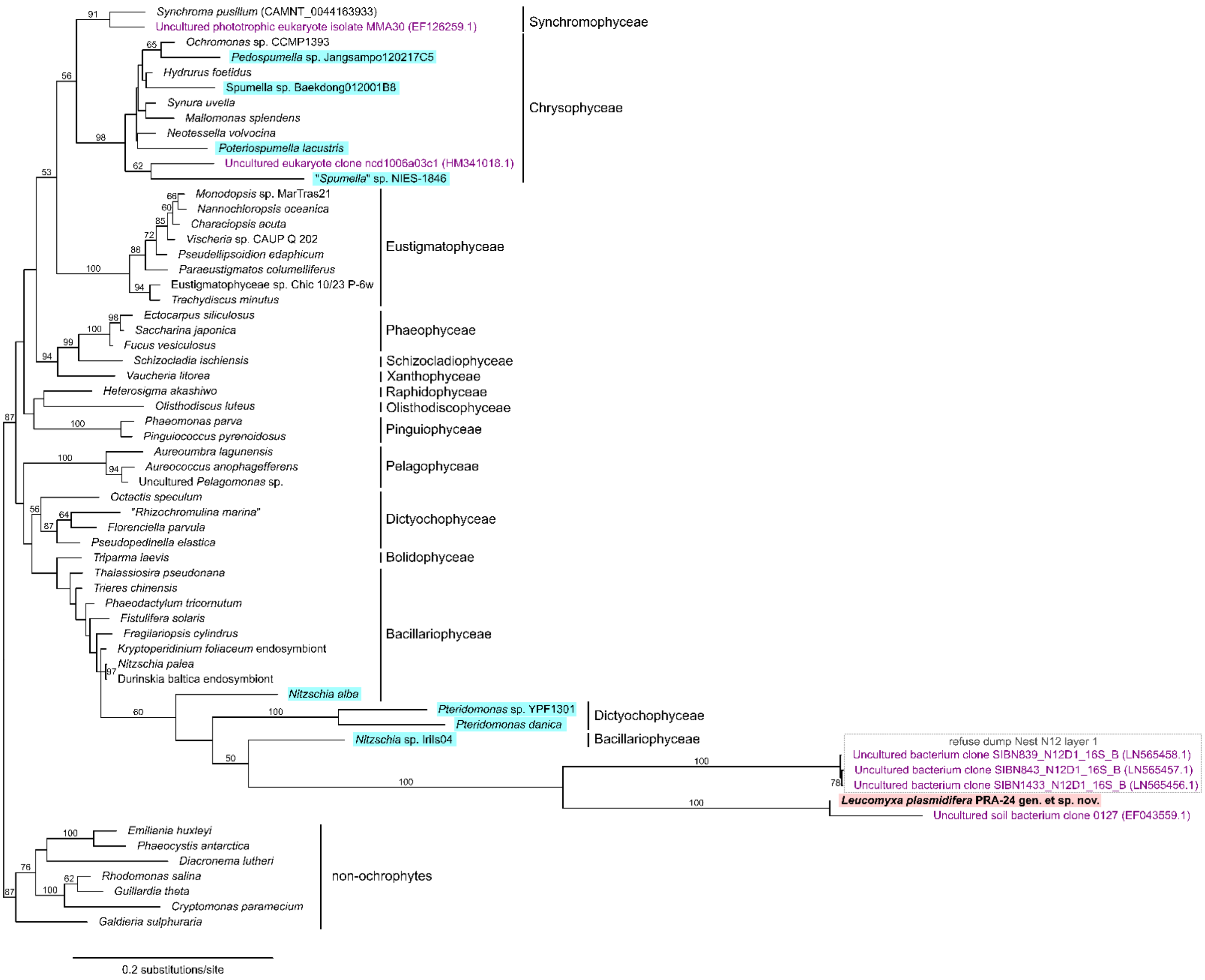


**Figure S4.** Maximum likelihood tree inferred from plastidial 16S rRNA sequences. The tree was inferred from a multiple sequence alignment of 1,236 nucleotide positions (after trimming) using IQ-TREE multicore version 2.0.3 and the substitution model GTR+F+R4. Bootstrap support values (calculated from 100 nonparametric bootstrap replications) are shown when ≥50. Non-photosynthetic taxa are highlighted with a coloured background. The sequences typed in violet correspond to environmental DNA clones from uncultivated organisms identified among best BLASTN hits in the NCBI non-redundant nucleotide sequence database queried with the 16S rRNA sequences from *Leucomyxa plasmidifera* PRA-24 or *Synchroma pusillum*. The group of three environmental sequences in a box was derived from the same environmental sample (“refuse dump Nest N12 layer 1”) as some of the environmental *Leuc. plasmidifera*-related 18S rRNA sequences included in the tree displayed in figure 2 (see there), indicating they may correspond to the same original organism. Sequences with no GenBank accession number indicated in the figure were extracted from the respective plastid genome sequences (for sources see supplementary electronic material, table S1). The *S. pusillum* sequence was extracted from a contig (ID provided at the tip label) in a transcriptome assembly available from this species. Note the paraphyly of diatoms (Bacillariophyceae) in the inferred tree due to an apparent misplacement of the long branches of the two *Pteridomonas* spp. (Dictyochophyceae) and the *Leuc. plasmidifera*-containing clade (expected to cluster together with *S. pusillum*).

```
                                                                          M··K··A··I··A··W··L··R·
PHD1_int cacacgtttgcgcgggaaataacggctagttcccttgcaattgaggagatactaataaagcagagatgaaggcgattgcttggttgctgagg
PHD1_dis cacacgtttgcgcgggaaataacggctagttcccttgcaattgaggagatactaataaagcagagatgaaggcgattgcttggttgctgagg
         ********************************************************************************************

           ·C··C··L··L··L··L··A··V··C··L··A··F··Q··P··D··A··A··P··I··S··A··R··R··R··A··P··V··K··P··G·
PHD1_int   tgctgcctgctgctgctggccgtatgcctcgccttccagccggacgcagcccccatctccgccagaagacgcgctcccgtgaagcccggc
PHD1_dis   tgctgcctgctgctgctggccgtatgcctcgccttccagccggacgcagcccccatctccgccagaagacgcgctcccgtgaagcccggc
           ******************************************************************************************

           ·Q··S··R··N··P··R··T··K··P··E··P··A··P··V··A··P··Q··L··D··T··Q··P··Q··I··K··P··K··Q··P··A·
PHD1_int   cagtccagaaacccgaggacaaagccagaacccgcacccgttgctcctcagctggacacgcagccgcaaatcaagcccaagcagccggcg
PHD1_dis   cagtccagaaacccgaggacaaagccagaacccgcacccgttgctcctcagctggacacgcagccgcaaatcaagcccaagcagccggcg
           ******************************************************************************************

           ·Q··Q··K··P··A··I··K··R··Q··P··V··I··R··H··V··R··N··N··H··G··S··N··P··V··D··L··W··V··V··G·
PHD1_int   caacagaagcctgcgatcaagcgtcagcccgtgattcggcacgtgcggaacaatcatggaagcaaccccgttgacctgtgggtcgtggga
PHD1_dis   caacagaagcctgcgatcaagcgtcagcctgtgattcggcacgtgcggaacaatcatggaagcaaccccgttgacctgtgggtcgtggga
           *****************************.************************************************************

           ·A··G··E··L··G··Q··R··V··I··K··I··W··K··E··L··H··P··R··S··V··V··V··A··E··T··L··S··R··D··R·
PHD1_int   gcaggcgagctcgggcaacgagtgatcaagatttggaaggagctgcacccgcgatccgtggtggttgcggagacgctgtcgcgtgaccgg
PHD1_dis   gcaggcgagctcgggcaatgagtgatcaagatttgaaaggagctgcacccgcgatccgtggtggttgcggagacgctgtcgcgtgaccgg
           ******************.****************.******************************************************

           ·H··R··F··L··K··P··L··G··V··L··C··S··L··R··E··K··R··S··E··R··D··A··R··I··A··S··N··V··V··F·
PHD1_int   catcgattcctcaagccgctcggcgtgctgtgcagcctgcgagagaagcggtcggaaagagacgcccgcatcgccagcaacgtcgtgttt
PHD1_dis   catcgattcctcaagccgctcggagtgctgtgcagcctgcgcgagaagcggtcggaaagagacgcccgcatcgccagcaacgtcgtgttt
           ***********************.*****************.************************************************

           ·A··A··P··P··T··G··T··L··D··Y··A··L··E··V··K··D··A··C··R··L··W··N··K··G··G··R··F··V··F··T·
PHD1_int   gcggcgccacccacgggcacactggactacgcgctggaagtgaaggacgcctgcaggctgtggaacaagggaggccgattcgtcttcacg
PHD1_dis   gcggcgccacccacgggcacactggactacgcgctggaagtgaaggacgcctgcaggctgtggaacaagggaggccggttcgtcttcacg
           *****************************************************************************.************

           ·S··S··T··R··N··V··L··P··D··K··K··G··V··I··S··E··T··S··P··M··N··A··S··D··V··A··L··A··E··N·
PHD1_int   tccagcactcgcaacgtgcttccggacaagaagggtgtcatctccgagacgtccccgatgaacgcctccgacgtggcgctggcggagaac
PHD1_dis   tccagcactcgcaacgtgcttccggacaggaagggtgtcatctctgagacgtccccgatgaacgccttcgacgtggcactggcggagaac
           ****************************.***************.**********************.*********.************

           ·Q··T··L··L··H··D··G··T··I··V··R··L··A··G··L··Y··N··L··K··R··G··P··H··A··K··W··L··K··D··G·
PHD1_int   caaactctgctgcatgacggcacgatcgtgaggctcgctggcctgtacaacctgaagcggggcccgcacgccaagtggctgaaggacggg
PHD1_dis   caaactctgctgcatgacggcacgatcgtgaggctcgctggcctgtacaacctgaagcggggcccgcacgccaagtggctgaaggacggg
           ******************************************************************************************

           ·H··V··A··G··H··P··Q··G··T··V··S··L··I··H··Y··D··D··A··A··A··A··V··V··K··A··L··L··H··G··Q·
PHD1_int   cacgtggcaggcccacccgcagggcaccgtctcgctcatccactacgacgacgcagcggcggcggtcgtcaaagcgctgttgcatggccaa
PHD1_dis   cacgtggcaggcccacccgcagggcaacgtctcgctcacccaccacgacgacgcagcggcggcggtggtcaaagcgctgttgcatggccaa
           **************************.***********.****.**********************.***********************

           ·A··N··A··T··Y··L··V··A··D··D··Q··A··L··T··R··K··Q··I··C··E··A··A··L··Q··H··P··R··F··C··E·
PHD1_int   gccaatgccacgtacttggttgccgacgaccaggcgttgacccggaagcagatctgcgaggcggcgctgcagcatccgcggttctgcgag
PHD1_dis   gccaatgccacgtacttggttgccgacgaccaggcgttgacccggaagcagatctgcgaggcggcgctgcagcatccgcggttctgcgag
           ******************************************************************************************

           ·K··S··M··P··T··F··D··S··S··A··P··K··V··K··K··S··C··N··T··T··W··T··R··K··Q··L··K··F··E··P·
PHD1_int   aagtcgatgccaacgttcgactcgagcgcgcccaaagtcaagaagtcctgcaacaccacctggacgcgcaagcagctcaagttcgagccc
PHD1_dis   cagtcgatgccaacgttcgactcgagcgcgcccaaagtcaagaagtcctgcacaccacctggacgcgcaagcagctcaagttcgagccc
           .***************************************************.*************************************

           ·V··Y··K··T··F··E··A··F··I··K··A··D··I··A··K··R··S··K··*
PHD1_int   gtgtacaaaacgttcgaggcgttcatcaaggcggacatcgccaagcgcagtaagtgatgtagaaactttgctgttgggttgtggcagatt
PHD1_dis   gtgtacaaagcgttcgaggcgttcatcaaggcggacatcgccaagcgcaataagtgatgtagaaactttgctgttgggttgtggcagatt
           *********.***************************************.****************************************

PHD1_int   ggaggtcggcctggttggacctgcgccccccttcgttctgccccttccacaaaaatcacagcacgtctgttgtgttctttgttgtagatgt
PHD1_dis   ggaggtcggcctggttggacctgcgccccccttcgttctgccccttccacaaaagtcacagcacgtctgttgtgttctttgttgtagatgt
           *****************************************************.************************************

PHD1_int   gttgtggtcttaatccctcgtgatttttcgacgaacgcataattttcccg
PHD1_dis   gttgtggtcttaatccctcgtgatttttcgacgaacgcataattttcccg
           **************************************************
```

**Figure S5.** Two versions of the PHD1 gene in *Leucomyxa plasmidifera* PRA-24. The figure shows a pairwise alignment of two different variants of PHD1 transcripts, each supported by multiple RNAseq reads mapped to the respective sequence present in the *Leuc. plasmidifera* transcriptome assembly obtained with the Trinity software (TRINITY_DN1429_c0_g1_i2). One variant (PHD_int) contains an intact coding sequence (its conceptual translation is included above the alignment), whereas the other (PHD_dis; identical to TRINITY_DN1429_c0_g1_i2), differs from the former by multiple substitutions, including synonymous (in yellow) and non-synonymous (in green); two substitutions each result in an in-frame termination codon (highlighted in red) disrupting the coding sequence.

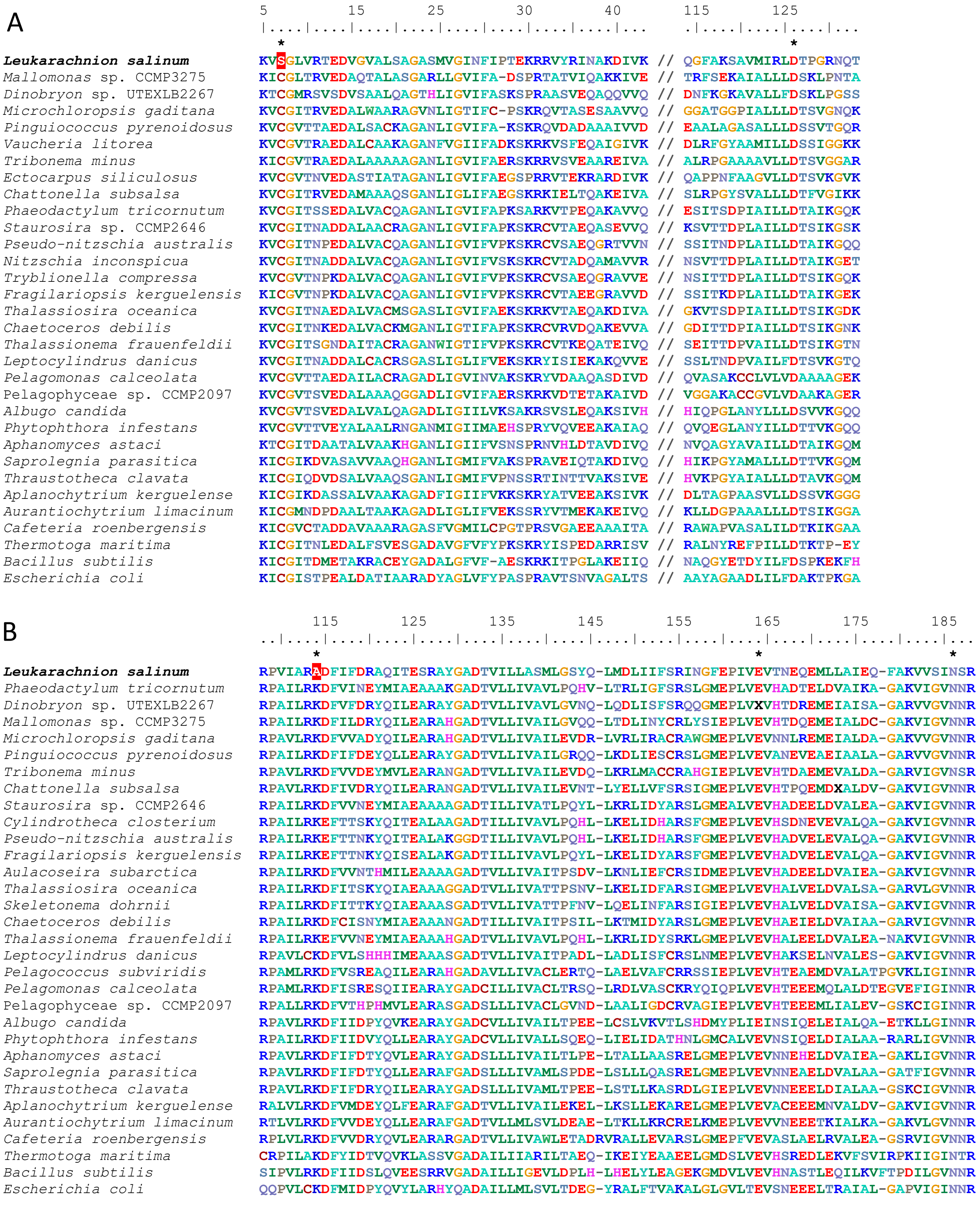


**Figure S6.** Mutations of catalytically critical residues in the TrpF-TrpC fusion protein from *Leucomyxa plasmidifera* PRA-24. The figure shows multiple sequence alignments of the regions containing residues established by experimental studies to be critical for catalytic activity of phosphoribosylanthranilate isomerase (TrpF; panel A) and indole-3-glycerol phosphate synthase (TrpC; panel B). These residues are indicated with the asterisks above the alignment. Residue numbering on the top corresponds to positions in the experimentally characterized TrpF protein from the bacterium *Thermotoga maritima* and the TrpC protein from the bacterium *Escherichia coli*. In addition to several reference bacterial sequences (bottom) and the corresponding regions from the *Leuc. plasmidifera* fusion protein (top), the alignments include selected TrpF and TrpC sequences from various stramenopiles, including chrysophytes closely related to *Leuc. plasmidifera* (immediately beneath the *Leuc. plasmidifera* sequence). Note that in contrast to all other sequences included in the alignment, the TrpF and the TrpC region of the *Leuc. plasmidifera* fusion protein each has a mutation of one of the invariant residues (highlighted in red) directly implicated in catalysis, suggesting loss (or at least reduction) of the catalytic activity.

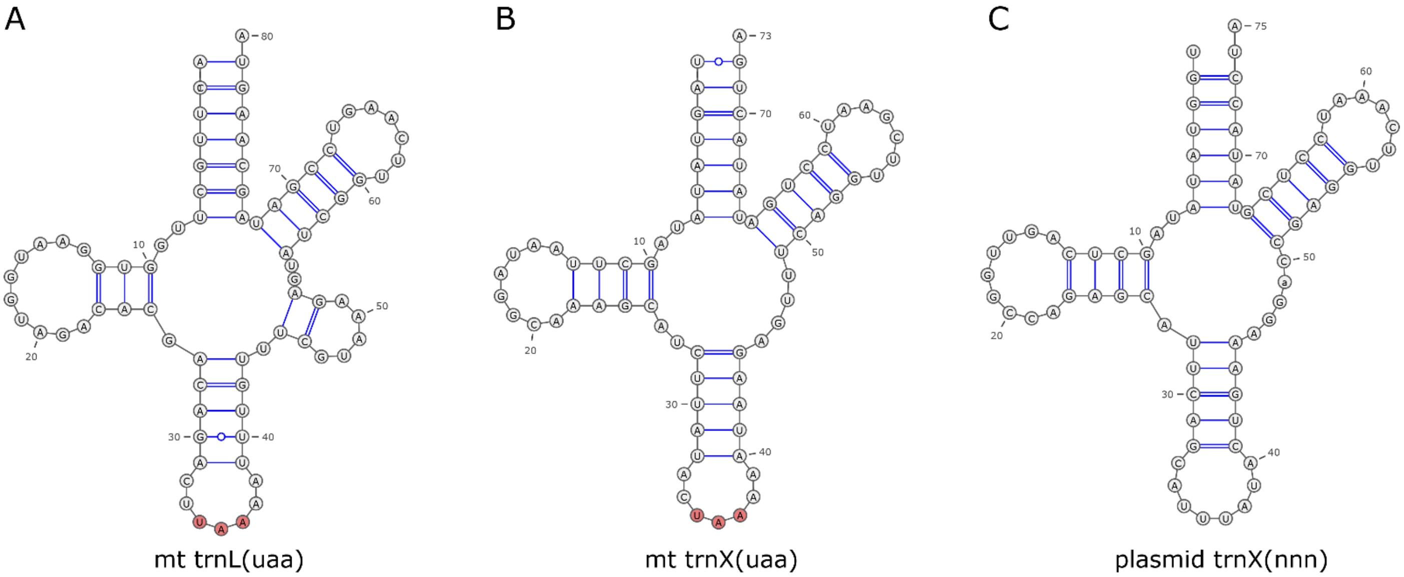


**Figure S7.** Unusual tRNAs specified by the *Leucomyxa plasmidifera* PRA-24 mitochondrial genome and its novel mitochondrial plasmid. (A) The standard trnL(uaa), a product of the *trnL(taa)* gene in the main part of the *Leuc. plasmidifera* mitogenome. Note the presence of a variable arm, a characteristic expected for a tRNA charged with leucine. (B) A tRNA specified by a gene, annotated as *trnX(taa)*, located in the long insertion in the first *cob* intron in the *Leuc. plasmidifera* mitogenome. Despite the presence of the anticodon UAA, the tRNA does not bear characteristics of leucine specific isoacceptors. (C) A tRNA specified by the single tRNA gene, annotated as *trnX(nnn)*, present in the *Leuc. plasmidifera* plasmid pLPM. Note that the anticodon loop is expanded by one nucleotide (has eight as opposed to the standard seven nucleotides), making its anticodon identity (and functionality in general) uncertain. Furthermore, as in the case of the tRNA shown in panel B, its amino acid specificity is also unclear.

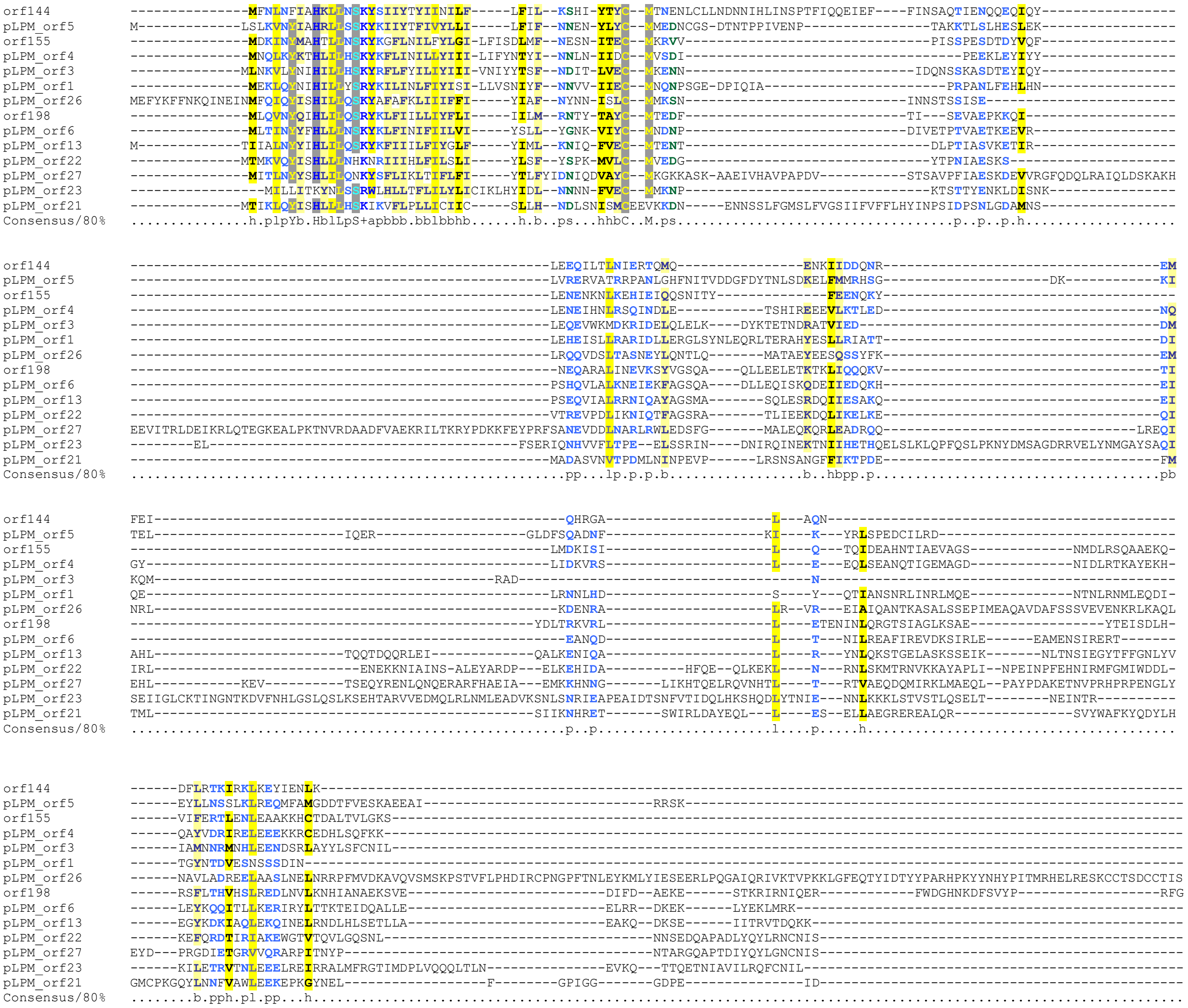

**Figure S8.** Novel protein family encoded by the *Leucomyxa plasmidifera* PRA-24 mitochondrial genome and its novel mitochondrial plasmid. Tree sequences included in the alignment are encoded by the mitogenome (orf144, orf155, and orf198), the remaining eleven (with the prefix “pLPM”) by the plasmid. The sequences were aligned using MAFFT version 7 (default settings) and further processed with CHROMA to highlight conserved positions. Note that only the N-terminal region is substantially conserved across the whole family.
